# Centromere inactivation during aging can be rescued in human cells

**DOI:** 10.1101/2023.12.30.573721

**Authors:** Sweta Sikder, Songjoon Baek, Truman McNeil, Yamini Dalal

**Affiliations:** Center for Cancer Research, National Cancer Institute/NIH, Bethesda, MD 20892; Saint Olaf College, 1520 St. Olaf Avenue Northfield, MN 55057

**Keywords:** Centromere inactivation, non-coding transcription, histone demethylase, aging, p53

## Abstract

Aging involves a range of genetic, epigenetic, and physiological alterations. A key characteristic of aged cells is the loss of global heterochromatin, accompanied by a reduction in canonical histone levels. In this study, we track the fate of centromeres during aging in human cells. Our findings reveal that the centromeric histone H3 variant CENP-A is downregulated in aged cells, in a p53-dependent manner. We observe repression of centromeric noncoding transcription through an epigenetic mechanism via recruitment of a lysine-specific demethylase 1 (LSD1/KDM1A) to centromeres. This suppression results in defective de novo CENP-A loading at aging centromeres. By dual inhibition of p53 and LSD1/KDM1A in aged cells, we mitigate the reduction in centromeric proteins and centromeric transcripts, leading to mitotic rejuvenation of these cells. These results offer insights into a novel mechanism for centromeric inactivation during aging and provide potential strategies to reactivate centromeres.

## Introduction

Cellular senescence was initially observed in the 1960s in human diploid cells cultured under laboratory conditions. These cells were found to have a maximum capacity for replication, beyond which they underwent a halt in their cell cycle progression, often referred to as the Hayflick limit ^1^. Cellular senescence is characterized by an arrest in either the G1 or G2 phase of the cell cycle. It is a state of irreversible growth arrest triggered by various mechanisms, such as DNA damage and telomere dysfunction. This state can be induced through multiple pathways, such as activation of oncogenes or loss of tumor suppressor functions, epigenetic alterations, or organelle damage ^2^. In addition to their irreversible cell growth characteristics, an important feature of senescent cells is the senescence-associated secretory phenotype (SASP). It involves the secretion and release of various cytokines, chemokines, growth factors, and proteases, which are secreted into the extracellular environment ^3,4^. Senescent cells release SASP factors that can negatively impact nearby cells and tissues, resulting in an accumulation of senescent cells and contributing to organismal aging ^5^. In recent years, research has focused on understanding how nuclear and chromatin changes trigger cellular senescence, thereby leading to organismal aging ^6^. It is now known that there is a rapid remodeling of the chromatin state characterized by alterations in histone modifications and DNA methylation patterns during aging in human cells ^7,8^. Deficiencies in key kinetochore components ^9^, such as BubR1, can contribute to aneuploidy and phenotypic changes associated with premature aging ^10^. However, whether centromeres, the chromosomal base upon which kinetochores are built ^11^, are altered during the aging process has not yet been elucidated. In this report, we ask a fundamental question in chromosome biology - what happens to centromeres during aging in humans?

Centromeres in humans have unique chromatin structure, including the presence of tandem arrays of approximately 171 base pairs of AT-rich α-satellite DNA ^12,13^ which are further organized into higher-order repeats ^14^. Most chromosomes (other than the human Y chromosome) contain a specific sequence motif known as the Centromere protein-B box within their higher-order repeats, which plays an essential role in recruiting CENP-B protein to the centromere ^15^. Centromere identity is primarily epigenetic and marked by a specific histone H3 variant CENP-A ^16^. CENP-A nucleosomes create a unique chromatin structure and are crucial for assembling functional kinetochore complexes ^17,18^. Along with 16 other proteins, CENP-A forms the constitutive centromere-associated network (CCAN) ^19^, with CENP-C playing a central role in facilitating interactions with both the Mis12 complex on the chromatin side ^20^. Depletion or reduced levels of CENP-A and CENP-C lead to cell division defects and chromosome mis-segregation ^21^ ^11^. Multiple factors, such as Mis16/RbAp46, the Mis18 complex, and HJURP ^22–24^, have been identified as playing crucial roles in facilitating the recruitment of CENP-A to centromeres. Traditionally considered transcriptionally inactive regions, it is now well accepted that centromeric transcription and transcripts are essential elements for the maintenance of centromeric epigenetic memory ^25–27^. In murine cells, the levels of these non-coding centromeric transcripts are tightly regulated during the cell cycle to maintain normal cellular function^28^.

An earlier work suggested a potential reduction of CENP-A levels in DNA damage-induced senescent fibroblasts, which may be regulated by the p53 pathway ^29^. However, this study had a limited understanding of the centromere function and alterations during aging. Here in this report across various ages of human donor tissues, we report a striking inverse relationship between age and CENP-A protein levels. To understand this phenomenon, we developed cellular senescence models to test how centromere specific proteins and centromeric transcription are perturbed during aging. Our findings show a progressive loss of canonical histones and redistribution of repressed chromatin domains. We observed a differential expression and distribution of histone variants and chromatin-associated proteins, including the loss of CENP-A and CENP-C. It also led to the suppression of centromeric transcription. We report a potential means to reactivate centromeric transcription in aged cells using a modulator of lysine-specific demethylase 1 and explore the role of CENP-B in regulating centromere function during aging. Together, these findings provide a plausible epigenetic mechanism for centromeric inactivation during aging. The implications extend to strategies for its reactivation, presenting a promising avenue for enhancing cellular health with advancing age.

## Results

### 1. Essential inner centromeric proteins CENP-A and CENP-C are downregulated in aging cells

To investigate the expression profiles of kinetochore-related genes and centromere components, we analyzed a comprehensive RNAseq dataset from the Genotype-Tissue Expression database (GTEx), encompassing various human tissue samples across different age groups. Transcriptomic data from kidney (n=64), lung (n=385), liver (n=158), colon (n=570), heart (n=626) and skin (n=1258) tissues were segregated into three distinct age clusters: young (20-39), middle (40-59) and old (60-79) (Figure 1A). The analysis revealed that most inner and outer kinetochore genes exhibited decreased expression in the older age group. Furthermore, the essential centromeric genes CENP-A and CENP-C displayed differential expression patterns with advancing age. Specifically, CENP-A and CENP-B were found to be downregulated in the older age group across most tissue types. In contrast, CENP-C demonstrated a mixed expression profile, with reduced levels observed particularly in kidney tissues. Building up from this analysis we first studied the endogenous expression of essential inner centromere proteins CENP-A/CENP-C in skin fibroblasts donated from healthy individuals of advancing age (n=9). Immunofluorescence analysis on interphase cells derived from these fibroblasts demonstrate that CENP-C protein levels significantly decline at 44 years of age and continue to remain suppressed in fibroblasts from older individuals. (Figure 1B). We also estimated the levels of H3K9me3, a heterochromatin mark, which tends to degrade with age ^30^. Consistent with the literature, we observed reduced levels of H3K9me3 in the nuclei of old fibroblasts (Figure 1A), and redistribution of the H3K9me3 at the periphery of the older nuclei (Figure 1A). These data are consistent with a potential reorganization of the global heterochromatin structure in aged cells. To further evaluate expression levels of centromere proteins we performed immunoblotting of whole cell extracts from these young and old fibroblasts. A comparable decrease in CENP-A protein levels with age, (1 year old versus 75 years old) as well as the degradation of Lamin B-1, the well-established marker for senescence was observed ^31^ (Figure S1B). Immunoblotting results for CENP-C also show a similar decline in 75-year-old fibroblasts versus the young age group of 26-year-old and 29-year-old. Interestingly, no alterations in the centromere-binding protein CENP-B were detected in the aged fibroblasts (Figure S1A and B). We next assessed whether the observed decrease in CENP-A/CENP-C proteins occurred at the centromeres of the chromosomes during metaphase. The growth rate of 96-year-old fibroblasts is very slow and it was a technical challenge to obtain enough metaphase spreads to quantitate the CENP-A signals. We therefore analyzed the essential kinetochore protein CENP-C, which is an accepted surrogate for CENP-A, and its closest binding partner at active centromeres, by immunofluorescence staining on metaphase chromosomes from young 29-year-old versus 44-year-old and 65-year-old fibroblasts. Consistent with the CENP-C staining of interphase cells, we observed reduced levels of CENP-C on the centromeres from both 44-year-old, as well as from 65-year-old skin fibroblasts as compared to 29-year-old (Figure 1C). Interestingly, chromosomes from the older fibroblasts appeared decompacted, indicating that the global chromosome structure may also be impaired during aging (Figure 1C). We also analyzed the CENP-B levels in the metaphase chromosomes and found no significant change with age (Figure 1D). Together, these results suggest that with an increase in age, there is a decrease in specific centromeric proteins CENP-A/CENP-C in skin fibroblasts. The rather dramatic decline of CENP-A and CENP-C protein levels (greater than 2-fold) suggests a potential weakening of centromeres as cells age (Figure 1A, Figure S1). To explore the specific effect of CENP-A downregulation on aging, we used a Locked Nucleotide Antisense Oligos (LNA-ASO) to specifically target CENP-A in the young 1 year old skin fibroblasts (Figure S2A). Knockdown of CENP-A significantly increased the CDKN1A/p21 senescent marker and led to degradation of Lamin B1 (Figure S3 and S2B). This indicates that loss of CENP-A could potentially contribute to a cellular senescent pathway in young skin fibroblasts.

**Figure 1:**
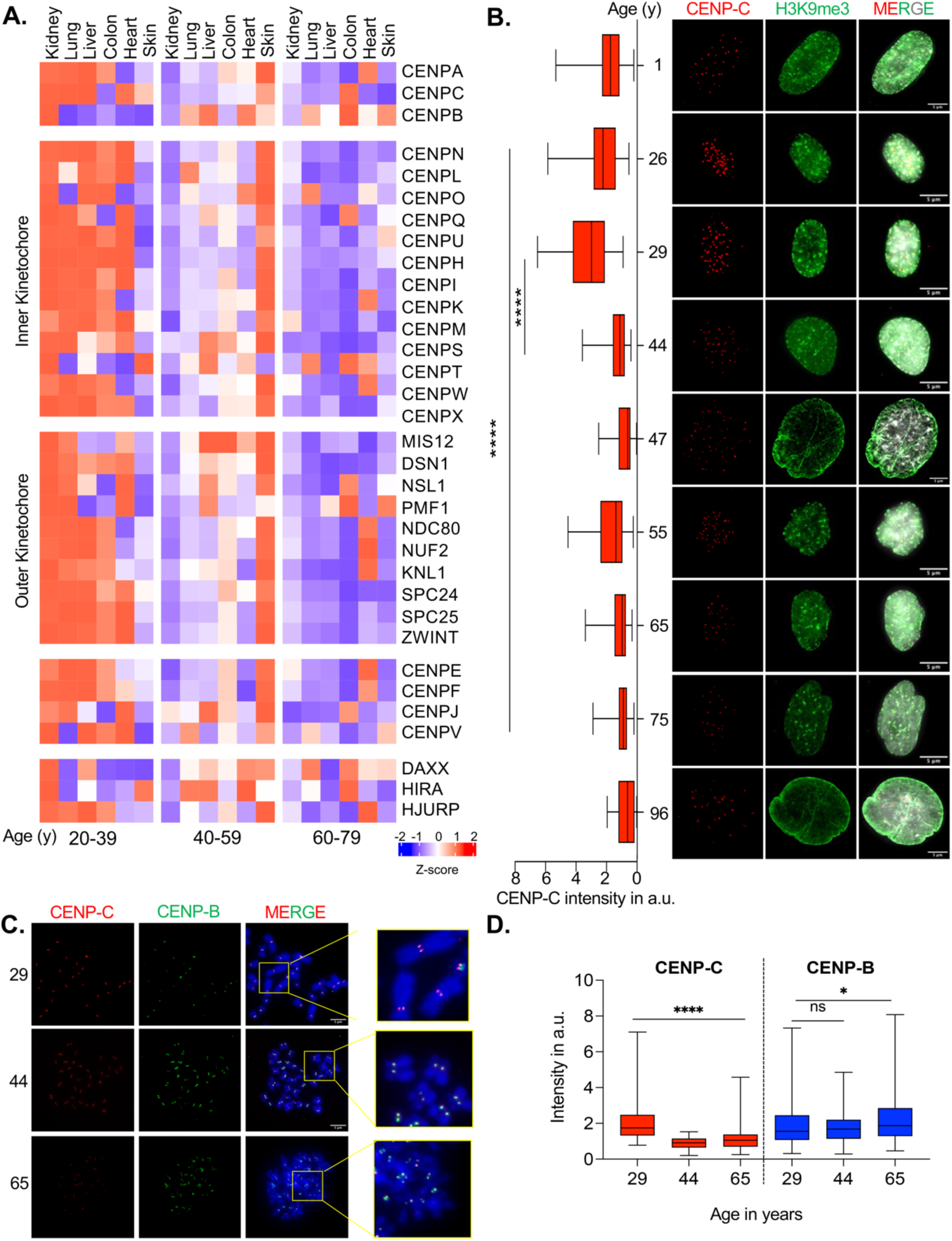
Centromere proteins decline in aged skin fibroblasts. (A) Heat map showing gene expression analysis of kinetochore related genes, and its chaperones of various human tissues as obtained from GTEx database. Each tissue dataset was classified into 3 age groups. All transcript data was normalized to the actin levels. Detailed analysis in the method section. Genes were classified according to their function in inner and outer kinetochore components, centromere genes, as well as chaperones. Most of the kinetochore related genes including centromere related genes were found to be downregulated in older age groups across different tissues (B) Representative IF image of CENP-C (in red) levels in different aged skin fibroblasts. H3K9me3 (in green) denotes the heterochromatin mark. Number of nuclei used for quantification denoted by n for each age is mentioned as follows: 1y (n=49), 26y (n=92), 29y (n=53), 44y (n=82), 47y (n=53), 55y (n=55), 65y (n=80), 75y (n=57), 96y (n=33) Scale bars as denoted. Single nuclei images imaged at 60X magnification except that for 1y, 47y, 96y which were imaged 100X magnification (B) Quantitation of fluorescence intensity of CENP-C (red) across different aged skin fibroblasts. CENP-C intensity diminishes in age 44 as compared to the young fibroblasts of age 29 year. (C) Representative metaphase chromosome IF-CENP-C (red) and CENP-B images of Young 29-year fibroblasts and age 44, 65-year skin fibroblast cells. The inlet shows zoomed-in images of chromosomes. Scale bars as denoted. (D) Fluorescence intensity of CENP-C and CENP-B scored on metaphase chromosomes from young as well as aged fibroblasts. Number of centromeric foci quantified for each group is as follows: 29y (n=129) 44y (n=153) 65y (n=105). Chromosomes from both 44 and 65-year-old skin fibroblasts exhibit reduced CENP-C levels. All data are shown as means ± SEM. **P < 0.001.

### 2. Centromeric protein expression and transcription are downregulated in induced aging

Working with chronologically aged skin fibroblasts in culture for an extended period presents significant challenges due to their limited growth potential. Therefore, we adopted previously established methods in the aging field, BJ skin fibroblast culture-based senescence system to study centromeric events in spatiotemporal detail (Experimental Procedures). We employed two currently established methods to induce cellular senescence: 1) BJ skin fibroblasts were either exposed to a low dose of doxorubicin (Aged^Dox^) for 24 hours followed by 10 days of incubation ^32,33^ or 2) continuously passaged (Aged^>PD^) until they reached the state of cellular senescence (population doubling time PD >80 for BJ) (Figure 2A). To verify that the doxorubicin treatment indeed induced senescence, we performed the beta-galactosidase assay and RNA-seq followed by a comprehensive differential gene expression analysis (Figure S5). Beta galactosidase assays demonstrate intense blue staining of the Doxorubicin treated cells, signifying the onset of senescence (Figure S5A). Additionally, gene expression analysis showed a notable rise in SASP-associated genes and a decrease in the lamin B1 gene. These properties are characteristic of senescent fibroblasts. (Figure S5C, S6A). Using this system, we next probed the levels of both centromeric proteins CENP-A and CENP-C. Across both the cellular senescence models (Aged^Dox^ and Aged^>PD^), CENP-A and CENP-C proteins were significantly downregulated (Figure 2B, D) when compared to the young fibroblasts (Figure 2B, D). We also examined the DNA damage levels in both the cellular senescence models. The doxorubicin treatment showed similar levels of **γ**-H2AX, a marker of DNA damage, as the senescent cells generated through continuous passaging (Figure 2C, D). This suggests that the decline in CENP-A/CENP-C levels is due to the induction of senescence (Figure 2D), and not solely because of increased DNA damage. We also checked the expression of H3.3 to verify whether the observed changes during aging are specific to the centromeric histone H3 variant CENP-A or extend to other histone H3 variants. Our immunoblotting analysis from whole cell extracts from young versus Aged^Dox^ cells validated the decrease in CENP-A expression in Aged^Dox^ cells while no notable alteration was detected in the H3.3 expression (Figure 2D, S4A). We also confirmed the decrease in CENP-A expression in doxorubicin-induced senescent lung fibroblasts (IMR90) and normal colon epithelial cells (Epicolon) (Figure S4B). This indicates that the loss of CENP-A during cellular senescence occurs across different cell types and varied tissue origin. Furthermore, we noted a significant increase in p53 protein expression (Figure S4A). This observation led us to consider the possibility of an inverse correlation between p53 and CENP-A expression in aged cells. To delve deeper into the pathways governing centromeric protein expression and cellular senescence, we performed a comparative analysis of gene expression utilizing RNA sequencing data obtained from both young and Aged^Dox^ cells (Figure S5A). A total of 2025 genes were globally perturbed, with a noteworthy proportion being repressed in aged cells (Figure S5B). Within the most downregulated genes, we identified CENP-A and BUB1 mRNA (Figure S6A), both crucial in the regulation of kinetochore and mitotic spindle assembly. Furthermore, pathway analysis of the RNAseq data unveiled dysregulation in genes associated with both mitotic and p53 pathways, suggesting possible crosstalk between these cellular pathways during aging (Figure S6B). To independently validate the RNA expression data for key targets, we performed qRT-PCR analysis of expression levels of specific senescence-related genes along with CENP-A mRNA. The qRT-PCR results demonstrated an upregulation of senescent-associated genes such as p21 (CDKN1A) and IL1A. Concurrently, the transcript levels of CENP-A and lamin B1 were notably downregulated in Aged^Dox^ BJ fibroblasts (Fig S7). Crucially, we also estimated the levels of centromeric transcripts through specific primers recognizing the hSATα RNA. Remarkably, the transcript levels of centromeric transcription were significantly downregulated in Aged^Dox^ BJ fibroblasts (Fig S7).

**Figure 2:**
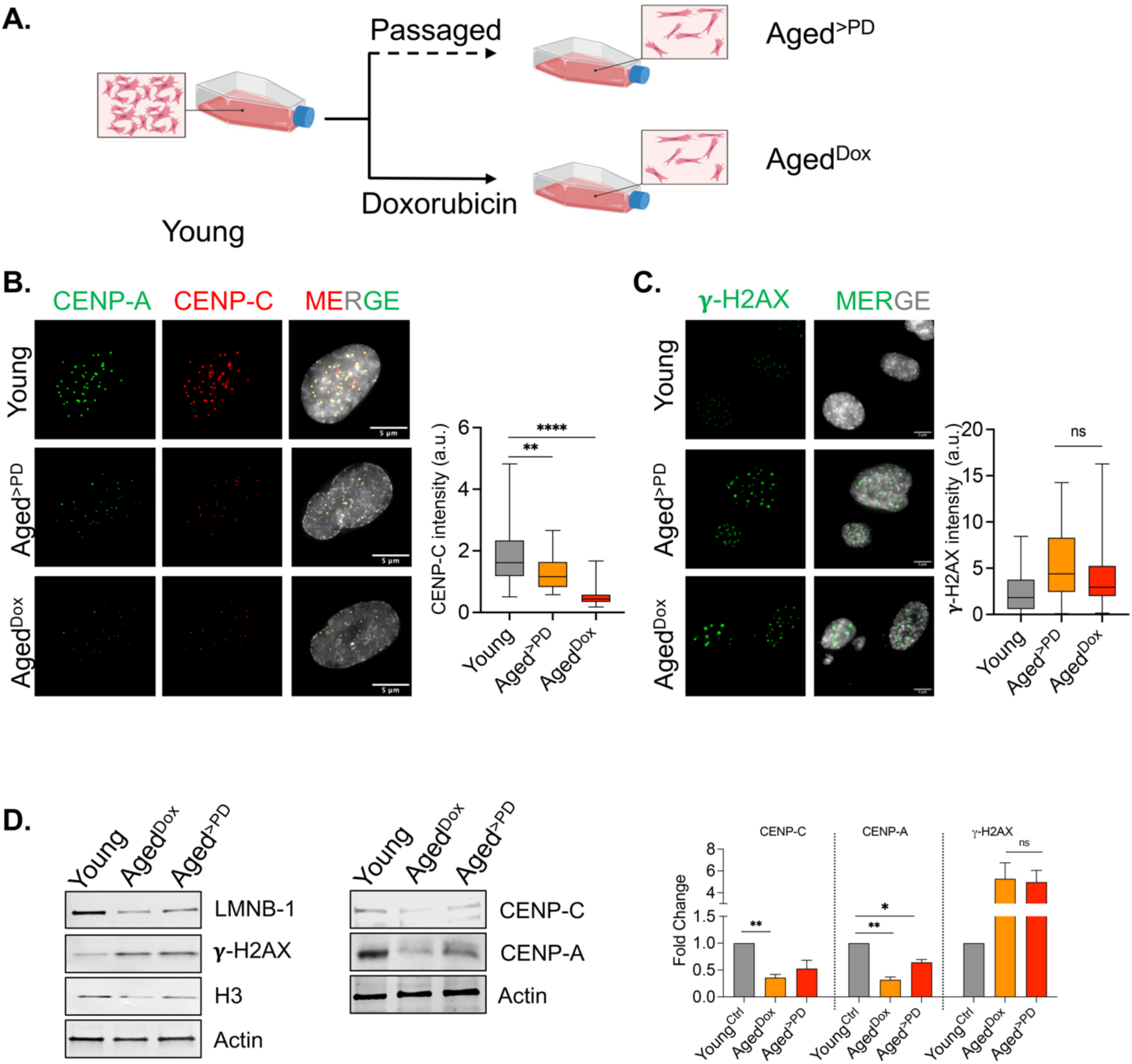
CENP-A/C expression decreases during doxorubicin-induced or replicative senescence of BJ skin fibroblasts. (A) Schematics of treatment conditions to induce senescence in vitro of BJ skin fibroblast cells. A low dose of DNA damage-inducing agent, doxorubicin (250nM) was administered to BJ cells for 24 hours. Following this treatment, the media was removed, and the cells were incubated for an additional 10 days to induce cellular senescence. Additionally, BJ cells were serially passaged until they reached greater population doubling times PD>80 and entered senescence. (B) Representative IF image of CENP-C (in red) levels and CENP-A (in green) in low population doubling time PD<10 BJ cells treated with DMSO as control as well as senescent cells as described above. Both replicative senescence (greater population doubling time) and doxorubicin-induced senescence reduced centromere protein levels. Quantitation of fluorescence intensity of CENP-C in young (n=81) and doxorubicin senescent BJ cells (n=79) and replicative senescent cells (n=50). CENP-C intensity significantly reduces upon Doxorubicin treatment-induced senescence (Aged^Dox^) as well as in BJ cells undergoing replicative senescence (Aged^>PD^). Number of nuclei denoted as n. (C) Representative IF image of **γ**-H2AX (in green) low population doubling time PD<10 BJ young cells treated with DMSO as control as well as senescent cells as described above. Both replicative senescence (greater population doubling time) and doxorubicin-induced senescence showed comparable levels of the DNA damage marker. Quantitation of fluorescence intensity of **γ**-H2AX in young (n=141) and senescent BJ cells. **γ**-H2AX intensity significantly increases upon Doxorubicin treatment-induced senescence (Aged^Dox^) (n=77) as well as in BJ cells undergoing replicative senescence (Aged>PD) (n=67). However, the difference of **γ**-H2AX levels in between both the senescent models are non-significant. (D) Representative immunoblot images of proteins isolated from BJ DMSO-treated young control cells doxorubicin-induced senescent cells, and aged passaged cells. CENP-A/C levels are reduced in senescent cells, but no significant difference is observed for histone **γ**-H2AX. Quantifications of immunoblots from 3 different biological replicates are represented in bar graphs. All data are shown as means ± SEM. **P < 0.001 and ****P < 0.0001.

Collectively, these findings imply that there is a decrease in centromeric CENP-A/C levels at both the transcriptional and translational levels across various cellular senescent cells indicating a universal phenomenon of centromere silencing upon aging ^34^. Additionally, aging might induce a possible transcriptionally repressive change at the centromere chromatin itself, thus shutting off centromeric transcription. This led us to investigate the global chromatin changes during aging and link them to the observed changes at the centromeres.

### 3. Chromatin fractionation analysis reveals changes in expression of centromeric proteins within differentially soluble chromatin fractions as cells age

We next isolated different transcriptionally accessible fractions of the chromatin - euchromatin and heterochromatin from young and Aged^Dox^ cells and examined protein expression changes. Aging is associated with a notable disruption in the global heterochromatin architecture within aged nuclei (Figure 1A) ^30^. However, the mechanisms responsible for the distinct patterns of compaction and decompaction in chromatin structure alteration during aging remain elusive. To delineate the protein expression pattern across the various chromatin states (euchromatin and heterochromatin), we adopted a widely used salt-differential chromatin fractionation protocol ^35^ followed by mass spectrometry analysis. Nuclei were obtained from both young and aged cells to extract chromatin. Through a process of gradual salt fractionation, it is possible to isolate specific chromatin states - lower salt concentration results in the isolation of euchromatin states or the more accessible fraction, while higher salt concentrations yield heterochromatin states (Figure 3A, S8). As expected, nuclear extracts from different salt fractions obtained from young cells revealed a clear separation of euchromatic and heterochromatic fractions, as indicated by the elevation of H3K9me3 levels in higher salt fractions (Figure S8B). In aged cells, there was a global reduction of H3K9me3 at higher salt fractions signifying the global loss of heterochromatin (Figure S8B). These nuclear fractions from both young and Aged^Dox^ cells were subjected to mass spectrometry (Figure 3A). Mass spectrometric analysis reveals a global perturbation of 1298 proteins in young vs aged cells. Identification of each of these proteins was confirmed by the detection of two or more unique peptides in the mass spectrometry data. Volcano plot depicts the differential expression of nuclear proteins both from euchromatin and heterochromatin fraction in young versus aged^dox^ cells (Figure 3B). A significant decrease of CENP-C peptide in aged fibroblasts compared to young nuclei was observed (Figure 3B). On the other hand, p53 binding protein TP53B and linker histone H1.0 levels were higher within aged fibroblasts (Figure 3B). In addition to the expression change, we also sought to examine the distribution patterns across various chromatin fractions of histone variants, including centromeric proteins and specific key lamin proteins that are known to be altered during cellular senescence. To examine this, we plotted the mz/ml ratio of each of these proteins across the increasing salt fractions (Figure 3C). Upon analyzing the distribution of histone variants and lamin proteins in different chromatin fractions, it was evident that most of these proteins were predominantly present in the low salt (0.5X PBS) euchromatin fraction in older cells (Figure 3C). We were also intrigued to find that, centromeric DNA-binding protein CENP-B is more abundant in the euchromatin fraction of aged cells. This observation also confirms our earlier finding that the CENP-B protein is present in aged fibroblasts (see Figure S1). Both CENP-B and CENP-C were detected in the euchromatin fraction in aged cells, in contrast to their predominant presence in the heterochromatin fraction in young cells (Figure 3C, S9). Thus, aging not only results in altered expression levels of centromeric proteins, but also a clear shift in their distribution across chromatin states. This comprehensive proteome analysis reveals that aging results in a distinct expression pattern of histone proteins, marked by a decrease in CENP-C protein levels. Intriguingly, TP53B protein and CENP-B were found to be expressed in aged cells, particularly within the more euchromatin-accessible fractions. Building upon these findings and our RNAseq data, we hypothesized that there could be a negative correlation between the p53 levels (Figure 2D) and CENP-A/CENP-C expression.

**Figure 3:**
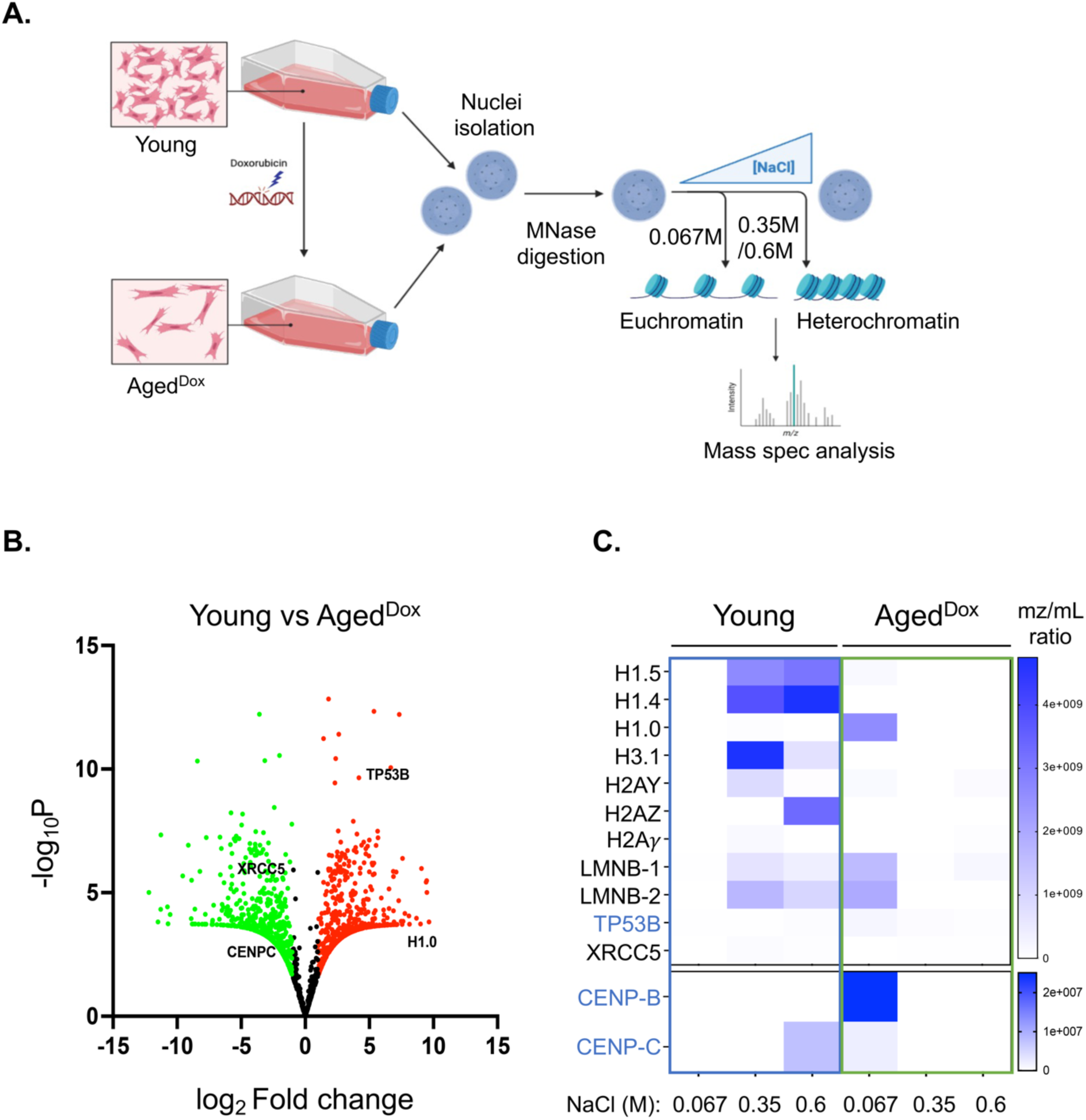
Proteomic analysis of different chromatin fractions from young and aged cells shows perturbation in centromeric protein levels. (A) Schematics of the experimental setup to analyze global proteome expression from young and doxorubicin-induced aged BJ cells. Nuclei isolated from both young and aged BJ cells were subjected to Micrococcal nuclease digestion (MNAse) followed by extraction in buffers with increasing salt concentrations. Lower salt concentration extracts euchromatin fractions while higher salt concentration isolates heterochromatin fractions. Both the chromatin fractions from young and aged cells were analyzed through mass spectrometry to determine differential protein expression. (B) Volcano plot of significant differentially expressed nuclear proteins in young vs aged cells. Nuclear proteins from young and aged fibroblasts from each chromatin fraction (euchromatin and heterochromatin) are represented together in the volcano plot. Proteins marked in green represent downregulated proteins whereas proteins marked in red represent upregulated ones. Proteins of interest are marked in black. CENP-C was detected among the downregulated proteins whereas TP53 and histone H1.0 were found to be upregulated. Biological replicates of n=2 represented. (C) Heat map representing the abundance of selected proteins, particularly histone variants across different chromatin fractions (increasing amount of salt). CENP-C was found to be downregulated while that of CENP-B was upregulated in aged BJ cells. Senescent markers like lamin B-1, and B-2 (LMNB-1/-2) were also shown to be downregulated in aged cells. Both centromeric proteins (CENP-B and CENP-C) were found to be expressed in the euchromatin fraction in aged cells as compared to the heterochromatin fraction of young cells.

### 4. Loss of p53 in aged cells upregulates CENP-A expression without rescuing mitotic defects

To test our hypothesis that p53 might play a regulatory role in CENP-A expression in aged cells, we knocked down p53 for 72 hours in Aged^Dox^ cells (Figure 4A) and analyzed CENP-A/C protein levels. Downregulation of p53 resulted in an upregulation of both CENP-A and CENP-C protein levels in aged cells (Figure 4B, C and S10). The former was expected based on previous work ^36^. The restoration of CENP-A expression was observed in 47-year-old skin fibroblasts (endogenous low level of CENP-A) following p53 silencing, indicating that p53 regulates CENP-A expression across various models of cellular senescence (Fig 4C, S10). Additionally, the decrease in p53 levels led to a reduction in p21 levels in the 47-year-old cells, suggesting a suppression of p21-mediated cellular senescence upon silencing of p53 (Fig S10). Based on this observation, we hypothesized that knocking down p53 in aged cells would suppress cellular senescence and the elevated levels of CENP-A would aid in ensuring correct cell division during mitosis. This prompted us to investigate Aged^Dox^ cells upon p53 knockdown during mitosis (Figure 4A). With a decrease in CENP-C protein, aging induces mitotic defects as compared to young cells (Figure 4B, C). Surprisingly, despite the observed increase in CENP-C foci, the mitotic defects in p53-silenced aged cells were not corrected (Figure 4B, D). We also monitored the aged cells upon p53 knockdown during interphase to look for markers of genome instability. We did indeed observe an increasing number of micronuclei formation (Fig S11). This led us to further investigate additional centromeric factors apart from CENP-A/CENP-C which might regulate correct mitotic processes. Our previous RT-qPCR results (Fig S7) showed that centromeric transcripts (hSATα RNA) was significantly repressed in Aged^Dox^ cells (Fig S7). It is now well accepted that noncoding transcription of repetitive elements at the human centromere plays a crucial role in the accurate deposition of CENP-A at the centromeres. Loss of such transcription results in loss of de novo CENP-A loading, and results in defective chromosomal segregation ^25^. Therefore, following p53 knockdown in Aged^Dox^ cells we next examined the levels of centromeric transcripts (Figure 4E). Silencing of p53 did not relieve the repression of centromeric transcription (Figure 4E). Thus, we speculated that the observed mitotic defects in cells with reduced p53 levels might be linked to suppression of centromeric transcription.

**Figure 4:**
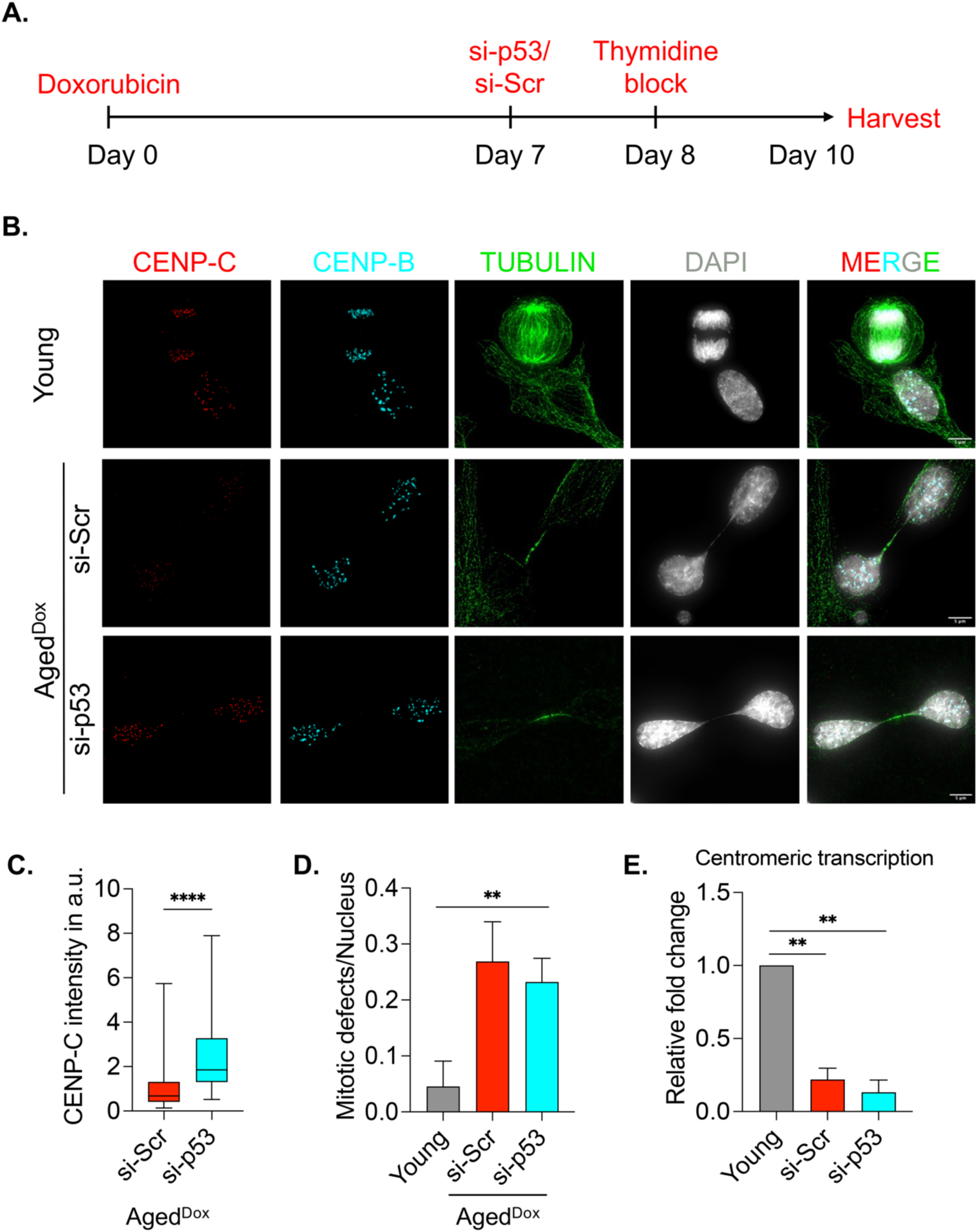
p53 silencing in aged cells harbor mitotic defects despite upregulation of CENP-A protein. (A) Schematic of treatment conditions and time scale of double thymidine block in both scrambled and p53 silenced doxorubicin treated senescent BJ cells. Thymidine was added to block cells at the G1/S phase and then released subsequently to analyze nuclei undergoing mitosis. (B) Representative IF images of cells undergoing mitosis in young control as well as aged scrambled and p53 silenced cells. Synchronized cells were immuno-stained for αTubulin (green), CENP-C (red), and CENP-C (cyan), and images captured and categorized as normal mitosis or defective mitosis (mitosis with lagging chromosomes, cytoplasmic bridges, multipolar spindles). Both scrambled and p53 silenced aged cells show extensive formation of nuclear cytoplasmic bridges resulting in segregation defects. (C) Quantification of CENP-C intensity in aged p53 silenced cells. Knockdown of p53 induces CENP-C expression in doxorubicin-induced senescent BJ fibroblasts. (D) Quantitation of mitotic defects in Aged^Dox^ cells (n=93) compared to young BJ cells (n=61), which persisted in p53 Aged^Dox^ knockdown cells. (n=142). (E) qRT PCR results showing centromeric transcript level using specific primers for hSATα transcripts in young and Aged^Dox^ cells transfected with specific siRNA (150nM) targeting p53 gene. Centromeric transcription is reduced significantly in aged cells. Its level does not change in aged cells upon p53 knockdown. Data from 3 biological replicates represented. All data are shown as means ± SEM. **P < 0.001.

### 5. Centromeric transcription is epigenetically repressed in aged fibroblasts

Centromeric transcription plays a crucial role in the incorporation of CENP-A and the stability of kinetochores ^25,26^. Existing research indicates that specific histone modifications, particularly H3K4me2, are vital in regulating centromeric transcription ^37,38^. To explore the potential epigenetic factors affecting centromeric transcription, we first examined the occupancy of heterochromatin mark at the centromeric DNA in aging cells. Chromatin immunoprecipitation, using an H3K9me3-specific antibody followed by RT-qPCR for centromere-specific hSATα DNA (α-sat) was performed in both young and Aged^Dox^ cells. Primers specific to the pericentromeric specific satellite hSATII DNA (sat II) chromatin region was also used as a control genomic region flanking the centromeric DNA. In Aged^Dox^ cells, both centromeric and pericentromeric DNA showed increased occupancy of H3K9me3 as compared to young cells, signifying the heterochromatinization of the centromeric region (Figure 5A). In addition to the repressive mark accumulation, we also analyzed the occupancy of the H3K4me2 transcriptional activation mark within the centromeric satellite DNA. We observed a notable decrease in the H3K4me2 levels, at centromeric hSATα DNA (Figure 5B) upon aging, but not at the pericentromeric satellite II DNA (Figure 5B). This synergistic reduction of H3K4me2 levels specifically at the centromeric DNA might explain the reduced expression of centromeric hSATα RNA in aged cells. The demethylation of H3K4me2 nucleosomes might be crucial for regulating centromeric transcription in aged cells. To test this hypothesis, we wanted to specifically inhibit lysine demethylase 1A (KDM1A/LSD-1) specifically, as it acts as a demethylase for the H3K4me2 nucleosomes at the centromeres ^39,40^. The small molecule inhibitor 2-PCPA specifically targets lysine demethylase 1A (KDM1A/LSD1) and inhibits KDM1A catalytic activity ^41^. We subjected Aged^Dox^ to 2-PCPA, which restored H3K4me2 levels (Figure 5C, S12A). Interestingly, the 2-PCPA treatment in Aged^Dox^ cells led to the upregulation of hSATα RNA expression as compared to the untreated control cells (Figure 5D). However, in contrast to Aged^Dox^ cells the LSD1i (2-PCPA) did not impact centromeric transcription in young cells (Figure 5D). This data suggests that the lysine demethylase 1A might regulate hSATα RNA expression specifically in aged cells. To confirm this phenomenon of centromeric transcription mediated by LSD1 in old fibroblasts, we exposed 47-year-old skin fibroblasts to 2-PCPA (Figure S12B). The inhibition of LSD1 resulted in increased expression of hSATα RNA in the 47-year-old fibroblasts compared to the untreated control. However, the administration of LSD1 inhibitor (2-PCPA) did not affect centromeric transcription in 1-year-old fibroblasts (Figure S12B) validating the regulation specific to aged cells. We next wanted to check if 2-PCPA could rescue centromeric transcription in p53-silenced Aged^Dox^ cells. Indeed, LSD1 inhibition showed similar rescue of hSATα RNA expression in p53 silenced aged fibroblasts (Figure 5E). Thus, this double repression of p53 and LSD1 in Aged^Dox^ cells established a system in aged cells where both CENP-A and centromeric transcription could be rescued. These intriguing findings made us curious to test whether the restoration of *both* CENP-A and hSATα RNA expression in aged cells could potentially mitigate mitotic defects.

**Figure 5:**
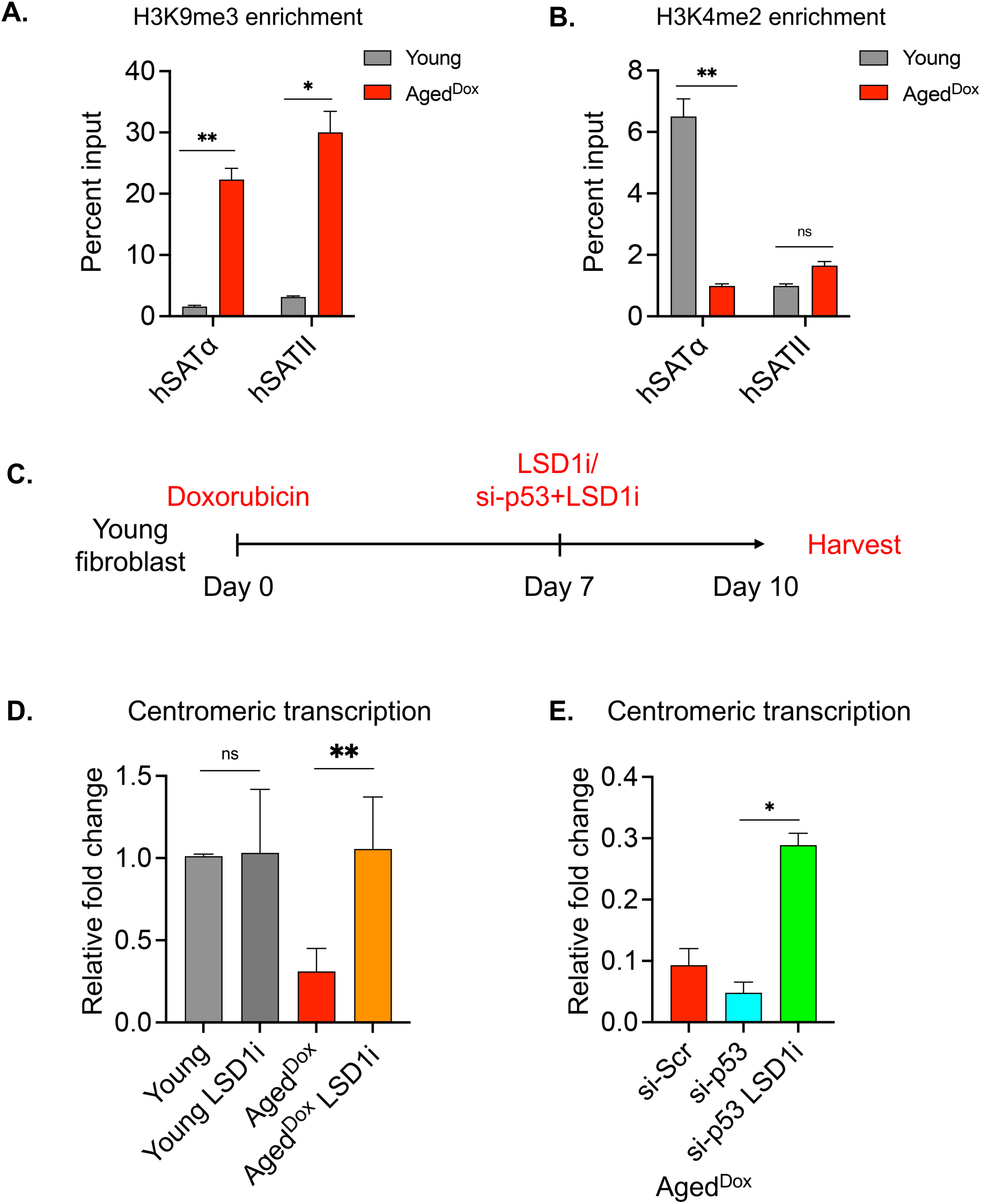
Centromeric transcription is regulated by H3K4me2 in aged BJ cells. Chromatin immunoprecipitation with (A) H3K9me3 and (B) H3K4me2 antibodies followed by qRT PCR with specific primers to measure enrichment at centromeric α-sat (hSATα) DNA and at pericentromeric DNA (hSATII) in young and Aged^Dox^ cells. Heterochromatin mark H3K9me3 was enriched at centromeric and pericentromeric regions in aged^Dox^ cells. H3K4me2 was significantly reduced at the centromeric α-sat DNA but not at the pericentromeric region. (C) Schematic to represent the experimental setup. Following doxorubicin treatment (for induction of senescence), 60µM of 2-PCPA (LSD1/KDM1A inhibitor) was added, and cells were harvested after 72 hours. For siRNA-mediated knockdown of p53, co-treatment of siRNA and 2-PCPA was done following 72 hours of incubation. (D) qRT PCR results showing centromeric transcript level using specific primers for α-sat transcripts in young and aged^Dox^ cells upon 2-PCPA treatment. Lysine demethylase inhibitor (2-PCPA) rescued centromeric transcription in aged cells. Data from 3 biological replicates represented. (E) qRT PCR results for quantitation of centromeric transcripts upon LSD1 inhibition in p53 silenced aged cells. LSD1 inhibition rescued centromeric transcription in p53-silenced cells. Data from 2 biological replicates represented. All data are shown as means ± SEM. *p<0.05, **p < 0.001.

### 6. KDM1A/LSD1 inhibition induces growth and rescues genome instability in p53-silenced aged fibroblasts

To investigate whether restoration of centromeric function, achieved by restoring both CENP-A/CENP-C and centromeric transcription, recovered the cell division defects, we analyzed p53 silenced cells undergoing mitosis. The double repression of p53 and LSD1 led to a significant reduction in mitotic defects compared to p53 knockdown alone (Figure 6A, B). We also observed a reduction in micronuclei following treatment with 2-PCPA in p53 silenced Aged^Dox^ cells (Figure 6C). Hence, inhibiting the catalytic function of LSD1 in p53-depleted Aged^Dox^ cells had the simultaneous effect of reinstating both hSATα RNA (Figure 5E) and CENP-A expression, leading to reduction of mitotic abnormalities. This observation encouraged us to determine the growth rate of aged cells upon both p53 knockdown and LSD-1 inhibition. Cell growth was assessed by quantifying EdU staining, which selectively identifies cells in S-phase. Notably, a substantial rise in EdU labeling was shown in Aged^Dox^ cells following the inhibition of LSD-1/KDM1A, indicating a marked enhancement in the growth of Aged^Dox^ cells (Figure 6D). Furthermore, when both p53 and LSD-1 were inhibited, we noticed a 2-fold increase of EdU-positive cells in the Aged^Dox^ as compared to the Aged^Dox^ control cells. This growth pattern of Aged^Dox^ by the dual inhibition of p53 and LSD1 was comparable to that of young cells (Figure 6D). Thus, inhibiting LSD1 in p53-deficient cells facilitated proper cell division, consequently rejuvenating the proliferative potential of aged cells.

**Figure 6:**
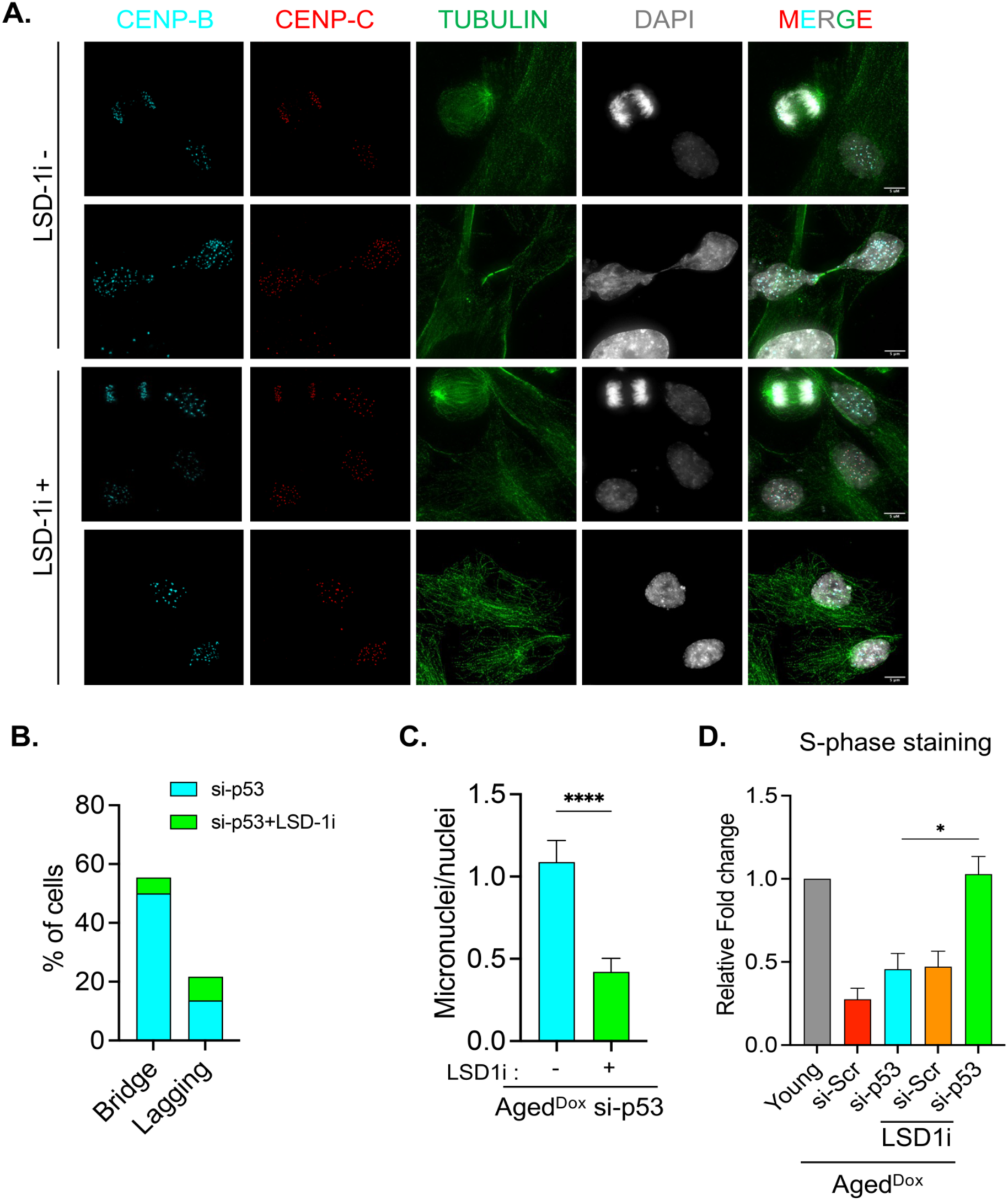
Lysine demethylase inhibition rescues mitotic defects in aged fibroblasts. (A) Representative IF images of cells undergoing mitosis in aged p53 silenced cells with and without LSD1 inhibition. Synchronized cells were immuno-stained for αTubulin (green), CENP-C (red), CENP-C (cyan), and images captured and categorized as normal mitosis or defective mitosis (mitosis with lagging chromosomes, nuclear-cytoplasmic bridges). LSD1 inhibition attenuated the mitotic defects observed in aged p53 deficient fibroblasts. (B) Quantitation of mitotic defects (percentage of both lagging chromosomes and cytoplasmic bridges) shows a significant reduction in p53 silenced aged fibroblasts treated with 2-PCPA for 72 hours. (C) Quantitation of micronuclei number in p53 silenced aged BJ fibroblasts upon 2-PCPA treatment. p53 silenced aged fibroblasts show a significant reduction in the micronuclei number upon LSD-1 inhibition. n>100. (D) Quantitation of EDU-positive cells from young and doxorubicin-aged fibroblasts upon treatment with LSD-1 inhibitor. Aged^Dox^ cells were transfected with scrambled or p53 siRNA to downregulate p53, followed by 2-PCPA (LSD1 inhibitor) treatment. Double knockdown conditions for both p53 and LSD1 showed maximum EDU positivity, signifying an abundance of replicating (S-phase) cells. Data represented of 3 biological replicates. All data are shown as means ± SEM. *P<0.05, **P < 0.001.

### 7. CENP-B regulates centromeric transcription by recruiting KDM1A to centromeres

The restoration of H3K4me2 and the rescue of centromeric function in senescent cells underscore the significance of recruiting KDM1A/LSD1 specifically at the centromeres during aging. We extended our investigation to determine the factor(s) that may aid the recruitment of lysine demethylase KDM1A at the centromeres. Our proteome analysis from aged cells (Figure 3) revealed that the centromere DNA binding protein CENP-B was stably expressed in the euchromatin fraction of aged cells. This prompted us to examine CENP-B expression levels in young and Aged^Dox^ cells. Our immunofluorescence data indicated no significant alteration in CENP-B expression in young versus Aged^Dox^ cells (Figure 7A and B). Expanding on the expression data, we postulated that CENP-B may recruit KDM1A/LSD-1 to centromeres, particularly within aged cells. This recruitment could potentially play a role in suppressing centromeric transcription during aging. To test this hypothesis, we employed a knockdown strategy of CENP-B using two specific antisense oligonucleotides in aged cells (Figure 7C). In line with our hypothesis, CENP-B knockdown significantly increased hSATα RNA expression in Aged^Dox^ cells (Figure 7D). Likewise, an increase in centromeric transcription was also observed upon CENP-B knockdown in skin fibroblasts derived from individuals aged 47 years old (Figure S13A). Furthermore, we aimed to determine if this increase in centromeric transcription is affected by the restoration of H3K4me2 at the aging centromeric DNA. Next, we examined the occupancy of H3K4me2 at the centromeric DNA in Aged^Dox^ cells following CENP-B depletion. ChIP-qPCR analysis showed significant enrichment of H3K4me2 upon CENP-B knockdown specifically at the centromeric DNA of Aged^Dox^ cells (Fig S13B). This observation suggests that CENP-B may epigenetically suppress centromeric transcription via the recruitment of KDM1A/LSD1. When CENP-B is knocked down, it disrupts LSD1 recruitment, resulting in the impaired demethylation of H3K4me2 nucleosomes. To further validate this, we expressed both Flag-CENP-B and HA-LSD-1 in both young and aged^dox^ cells. We fixed the cells 72 hours post transfection and performed co immunofluorescence using anti-mouse-flag tagged antibody (for CENP-B detection) and anti-rabbit-HA antibody (for HA-LSD-1 detection) (Figure 7E). Our co-immunofluorescence quantification data show greater percentage of colocalization of CENP-B with LSD-1 in aged cells as compared to young cells (Figure 7F). This suggests a greater interaction of CENP-B and LSD-1 in aged cells thus suggesting a plausible mechanism of centromeric inactivation through the CENP-B-LSD1 axis during aging.

**Figure 7:**
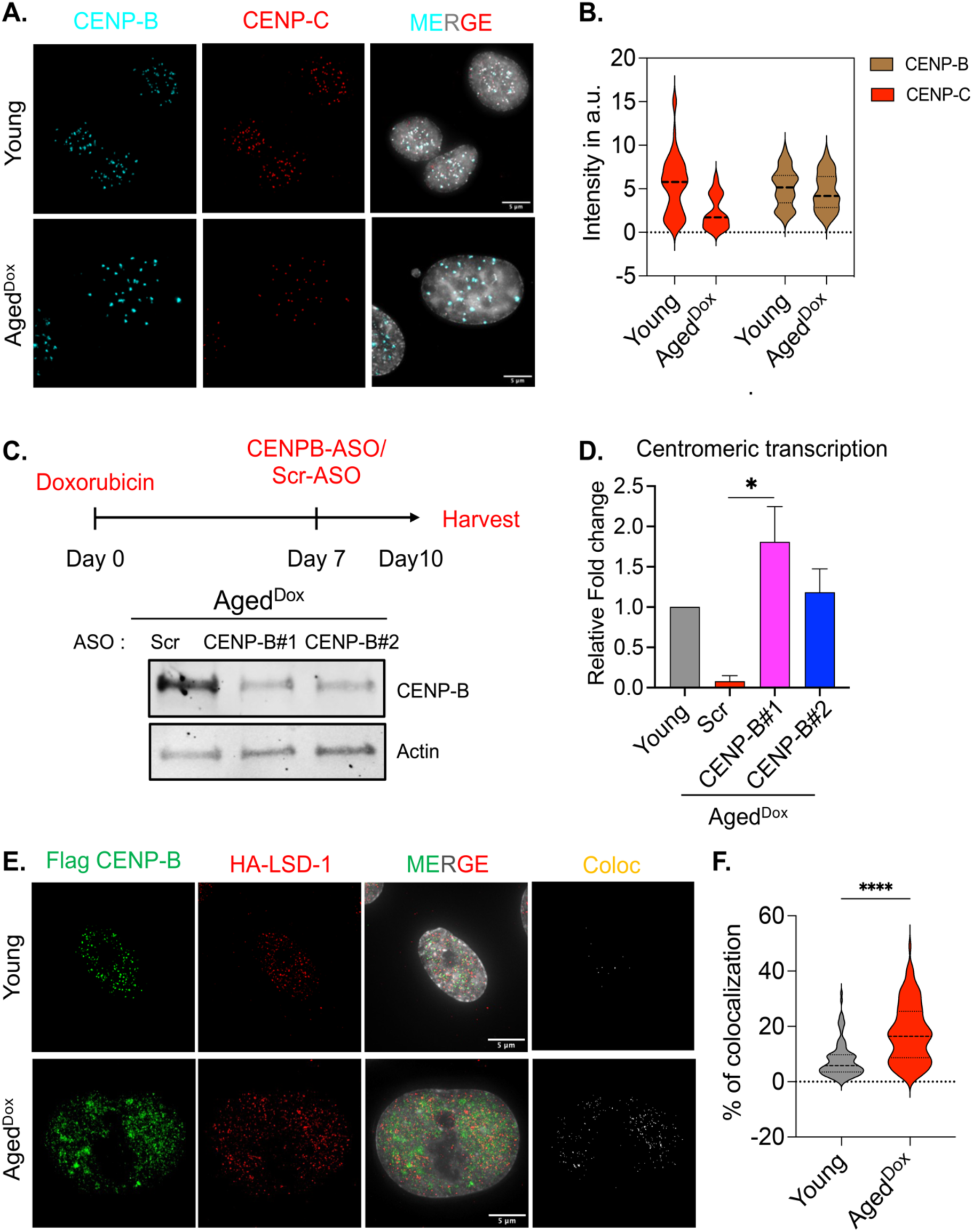
CENP-B epigenetically regulates centromeric transcription in aged fibroblasts. (A) Representative IF image of CENP-B (green) levels in young and doxorubicin-aged BJ fibroblasts. CENP-C is represented in red. Scale bars as denoted. (B) Quantitation of fluorescence intensity of CENP-B in young and Aged^Dox^ fibroblasts shows no significant change in its expression upon aging. (C) The upper panel represents the schematic of the transfection of ASOs targeting CENP-B in aged BJ fibroblasts. Lower panel: representative immunoblot image for estimating CENP-B protein levels upon transfection of ASOs in aged BJ fibroblasts. Both ASOs at 3µM showed a significant reduction in the CENP-B levels after 72 hours of transfection. (D) Quantitative real-time PCR for centromeric transcripts using hSATα primers upon CENP-B knockdown. CENP-B depletion results in significant upregulation of hSATα RNA in aged BJ fibroblasts. Data represented of 3 biological replicates. (E) Representative IF images of BJ Young and aged^dox^ cells expressing HA-LSD-1, Flag-CENP-B, post 72 hours of transfection. Co immunofluorescence with anti-mouse Flag antibody (green), anti-rabbit HA (red) antibody to show co localization of CENP-B and LSD-1. Scale bar as denoted. (F) Quantitation of colocalized points (white), shows greater co-localization of LSD-1 with CENP-B in aged^dox^ cells (n=114), as compared to young cells (n=113).

### 8. Centromeric inactivation is underpinned by epigenetic changes in centromere transcription in aging human tissues

We finally wanted to extend these conceptual findings of the epigenetic basis for centromeric inactivation, in human donor derived aging samples. To achieve this, we obtained and examined human lung and kidney donor tissues of varying ages. We analyzed the expression levels of centromeric proteins CENP-A and CENP-B, and centromeric hSATα RNA expression from these tissues. To analyze both protein and RNA expression, each tissue sample was divided into two fractions, enabling both protein and RNA analyses. Our findings indicate a reduction in centromeric transcripts as individuals age in both lung and kidney tissues (Fig 8A, C). It is worth noting that while the trend of declining transcript levels held across both tissue types, there were variations in the extent of fold change observed. Lung tissues exhibited a more than 2-fold reduction in hSATα RNA levels around the mid-age group (30-60 years). In contrast, kidney tissues demonstrated a sustained expression of hSATα RNA in the mid-age group, with a significant decline of > 2-fold observed only in the older-age group (>60 years) as compared to the young control samples (1–30-year-old). This intriguing variability underscores the heterogeneous nature of tissue aging, which is influenced by their distinct origins and characteristics. In line with our protein expression data for centromeric proteins in aging fibroblasts (Figure 1A, 2A), we also observed a distinct reduction in the expression of CENP-A in both aged lung and kidney tissues (as shown in Figures 8B and 8D). We also observed an increase in CENP-B expression upon aging in kidney tissues with advancing age (Figure 8B). As previously observed in the context of aged fibroblasts, these data suggest a potentially antagonistic role of CENP-B in regulating centromeric transcription.

**Figure 8:**
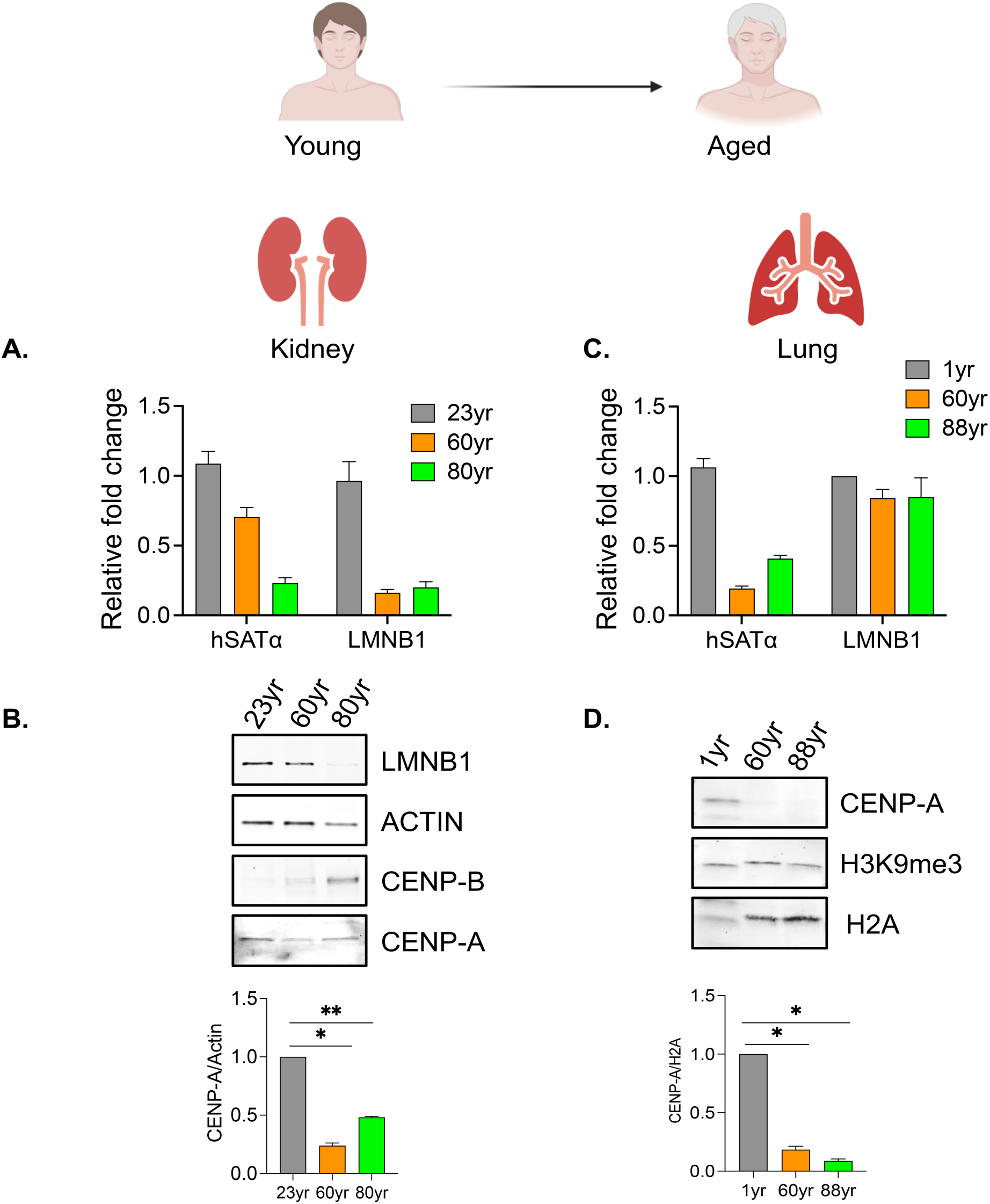
CENP-A and centromeric transcription is downregulated in aged human tissues. (A) Quantitative real-time PCR for centromeric transcripts hsat-α in different aged human kidney tissues. LMNB1 signifies lamin B1 transcripts. Gradual depletion of centromeric transcripts with age in kidney tissues are observed. Data represented of 3 replicates. (B) Representative immunoblot image for whole cell extracts of different aged kidney tissues show downregulation of CENP-A upon aging (C) Quantitative real-time PCR for centromeric transcripts hsat-α in different aged human lung tissues. LMNB1 signifies lamin B1 transcripts. Centromeric transcripts go down with age in kidney tissues. Data represented of 3 replicates. (D) Representative immunoblot image for whole cell extracts of different aged lung tissues show downregulation of CENP-A upon aging.

These observations lead us to conclude that the phenomenon of centromeric inactivation is consistently observed across various contexts in aging models, including chronologically aged fibroblasts, chemically induced senescent cells, as well as human-donor derived aged tissues.

## Discussion

In most multicellular eukaryotic organisms, aging is an inevitable process. It is often caused by the ongoing accumulation of DNA damage within cells ^42,43^. This leads to cellular arrest and eventual senescence. Centromeres represent tightly regulated genomic regions crucial for kinetochore attachment, and they undergo notable transformations during the cell cycle ^44^. While telomere attrition ^45^ is a well-acknowledged marker for aging, current understanding of the centromere remains surprisingly limited in the context of aging. A previous study has shown a higher order chromatin reorganization in centromeric DNA upon cellular senescence^46^, however the mechanistic insights still remain to be deciphered. This work aims to fill that fundamental gap in our knowledge.

Our data indicates a significant decline in the levels of the centromeric histone H3 variant CENP-A in cells and fibroblasts obtained from individuals spanning various ages, as well as in aged tissues (Figure1,2,8). We also mined a large human transcriptomic dataset obtained from different ages of 6 different human tissues to analyze the gene expression data of kinetochore and centromere related genes. Our data indicates a widespread downregulation of numerous kinetochore-associated genes (Figure 1A) in aged human tissues, which likely contributes to the mitotic abnormalities and impaired cell cycle progression observed in older individuals. Additionally, we also observed the downregulation of essential kinetochore protein CENP-C/A levels in the mitotic chromosomes of aged skin fibroblasts (Figure 1C, S1B). This decline of specific centromeric proteins was observed exclusively in aging tissues and cells, with no such decline noted in quiescent cells (data not shown). This reveals a distinct molecular phenomenon occurring at the centromeres as cells stop dividing and irreversibly exit the cell cycle. CENP-A/C is downregulated due to the increased levels of p53 in aged cells (Fig 4C, S10). The increase in p53 protein levels might be due to the accumulation of DNA damage during cellular senescence ^47,48^. The DNA damage response initiates the p53– p21-mediated senescence pathway ^49^. p21 acts as an inhibitor for cyclin-dependent kinase complexes, inhibiting cell cycle progression by thwarting DREAM (dimerization partner, RB-like, E2F, and multi-vulval class B) complex formation. The DREAM complex represses cell-cycle genes ^50^. Interestingly, our data shows when p53 was silenced in aged cells, there was a restoration of CENP-A/CENP-C expression along with the reduction of p21. However, these aged cells still exhibited mitotic defects. This is particularly intriguing, especially in the context of transformed or cancer cells, where non-functional wild-type p53 is often prevalent which might lead to CENP-A upregulation^51^ yet result in aneuploidy. The centromeric DNA though highly repetitive undergoes transcription to give rise to short noncoding RNAs ^27^. A previous study described that CENP-A-depleted cells experience abnormalities at centromeres, possibly arising from an imbalance in DNA recombination at these specific genomic regions ^52^. This increase in recombination could generate centromere erosion due to the shortening of repetitive DNA. To further investigate the centromeric factors that regulate the accurate segregation of chromosomes, we discovered that the centromeric transcription was silenced in aged cells and tissues (Fig 4E, Fig 8B, D). The repression of centromeric transcription (hSATα RNA) resulted through the accumulation of the H3K9me3 heterochromatin mark and the reduction of transcriptionally permissive H3K4me2 mark at the centromeric DNA in aged cells (Fig 5A, B). These results reinforce the possibility that the epigenetic loss of centromere function may be a significant contributing factor to accumulation of mitotic defects in aged cells.

We further elucidate a novel but distinct mechanism for regulating centromeric transcription to centromeric protein expression in aged cells. By inhibiting both p53 and KDM1A/LSD-1, we were able to rescue not only CENP-A but also hSATα RNA expression along with mitigating mitotic defects (Figure 5D, 6). Moreover, our study also emphasizes that centromeric function relies on both CENP-A expression and its accurate deposition at the centromeres through the centromeric noncoding hSATα RNA transcription. Our results also unveil a novel role for CENP-B in potentially regulating hSATα RNA expression in aged cells, possibly through recruiting KDM1A/LSD1 to the centromeric DNA (Figure 7C, D, E, S13). Additionally, we observed an increase in CENP-B and a decrease in CENP-A and centromeric transcription within kidney tissues of aged individuals. This suggests that the CENP-B-centromeric transcription axis may also be active in aging human tissues.

A striking new avenue for investigation that arises directly from our work is what happens at centromeres in other species when their cells undergo exit from the replicative cycle. It is also intriguing to speculate whether in aging cells, loss of targeted-CENP-A deposition that normally operates to faithfully restore centromere identity, also contributes inadvertently to off-target CENP-A/C deposition. If this is the case, one might predict a corollary to loss of correct centromere function is also gain of chromosome instability at ectopic centromere-like domains, which could contribute to age-related cancer progression. Pursuing this question in our lab is a logical current extension of the work presented in this report.

In sum, our study systematically unveils a novel mechanism for alleviating mitotic arrest in aging cells—by activating centromeres to restore cell cycle potential. The prospect of strategically reinstating epigenetic factors specifically at the centromeric DNA emerges as a promising avenue for targeted intervention in aging cells.

## Materials and Methods

### Cell culture

BJ human skin fibroblast cells were grown in a humidified 37°C incubator containing 5% CO_2_ in MEM medium containing glutathione and high concentrations of vitamins (catalog no. 11875093, Gibco, USA) supplemented with 10% fetal bovine serum (FBS) (catalog no. S11050H, Atlanta Biologicals, USA) and 1X penicillin-streptomycin solution (catalog no. 15140122, Gibco, USA). A total of 10 human fibroblast cultures, established from skin samples of Caucasian males with ages ranging spanning from 1yr to 96 yr old were acquired from cell biobanks as summarized (Key resource table). All donors were reported as “healthy”.

#### Doxorubicin-induced senescence

BJ skin fibroblast cells approximately 1X10^6^ were seeded on the previous day and were exposed to 250nM doxorubicin (Dox) (Doxorubicin hydrochloride) (Catalog number: D1515, Sigma -Aldrich) for 24h. On day 1, Dox was replaced with fresh media, and cells were followed until day 10 with regular media changes every 4 days. As a control, low population doubling time cells were exposed to DMSO for 24h and then monitored similarly for 10 days. BJ cells once confluent was passaged and seeded onto fresh flasks within the timespan of 10 days.

#### Lysine demethylase inhibition

Aged^Dox^ cells were treated with 60µM 2-PCPA for 72 hours. Untreated cells were kept as control. After 72 hours cells were harvested either for RNA isolation or for immunoblotting.

#### Senescence-associated b-galactosidase (SA-bGAL) assay

SA-bGAL activity was evaluated in the doxorubicin-induced senescent cells after 10 days using the Senescence Detection Kit (Catalogue number: ab6535, Abcam) as per manufacturer’s instructions. Briefly, cells were seeded into triplicates in 24-well plates, washed twice with 1X PBS, and then fixed with formalin for 15 min at room temperature. Next, cells were washed twice with 1 X PBS and incubated overnight at 37°C with b-gal substrate in an acidic buffer (pH 6.0). with an inverted microscope and photographed (Olympus BX41). The development of a perinuclear blue color was taken as an indicative of a senescent cell.

#### Cell proliferation assay (EdU staining)

EdU staining was conducted using Click-iT^TM^ EdU Flow cytometry kit (Invitrogen, Carlsbad, CA, USA) according to the manufacturer’s protocol. Cells were grown in an appropriate culture medium and were cultured and incubated with EdU reagent for 2 h. After the removal of EdU-containing media, cells were harvested and washed with 3 mL of 1% BSA in PBS. The pellet was then fixed with 100 μL of Click-iT® fixative and incubated for 15 minutes at room temperature, protected from light. After washing twice with 3% BSA in PBS, the slides were incubated with a Click-iT^TM^ reaction cocktail containing Click-iT^TM^ reaction buffer, CuSO4, Alexa FluorÒ594Azide, and reaction buffer additive for 30 min while protected from light. The cells were then washed with 3 mL of 1X Click-iT® saponin-based permeabilization and wash reagent. Cells were pelleted and 500 μL of 1X Click-iT® saponin-based permeabilization and wash reagent was added and then analyzed by flow cytometer.

### ASOs/si-RNAs and transfection

Locked nucleic acid (LNA) GapmeR ASOs were custom-designed and purchased from Qiagen, USA. A list of the sequences of LNA ASOs used in this study is provided in the key source table. siRNA specific for the TP53 gene was purchased from Dharmacon. Transfection of the LNA ASOs/siRNA was done using Neon™ Transfection System 100 µL Kit (catalog no. MPK10096, Thermo Fisher Scientific USA) following the manufacturer’s protocol. Briefly, the ASOs were added to 100 μl of Resuspension buffer R solution and incubated at room temperature for 5 min. In the Neon™ transfection system, the Neon™ Tube is filled with 3 mL of Electrolytic Buffer E2 so that the electrode on the side of the buffer is completely immersed in it. The resuspension buffer mix containing the cells with ASOs/siRNA was aspirated onto a 100ul with the recommended transfection program. After transfection, the cells were grown in antibiotic antibiotic-free MEM medium for 24 hours and harvested after 72 hours as per the experimental design.

### Nuclei isolation and MNAse digestion

Both young and aged cells were harvested and rinsed twice with ice-cold 1X PBS with 0.1% Tween. Nuclei were released with TM2 (20 mM Tris–HCl, pH: 8.0; 2 mM MgCl2) with 0.5% Nonidet P 40 Substitute (Sigma Cat #74385). Released nuclei were spun down at 1000 rpm for 10 min, rinsed once with fresh TM2 buffer, and re-suspended in 1 ml 0.1 M TE (10 mM Tris, 0.2 mM EDTA, 100 mM NaCl). Chromatin was either digested for 4 min for chromatin fractionation or 16 min min for ChIP-qPCR with 2.0 U MNase (Sigma Cat #N3755-500UN) in nuclei solubilized with ’ mL of 0.1 M TE (10 mM Tris, 0.2 mM EDTA, 100 mM NaCl) and supplemented with 1.5 mM CaCl_2_. MNase reaction was stopped by adding 10 mM EGTA, pH: 8.0 and pipetted up and down 3X to ensure even distribution and placed on ice for 5 mins. MNase-digested nuclei were spun down at 1500 rpm for 10 min.

### Chromatin Immunoprecipitation

Native ChIP experiments were performed on BJ Young and Doxorubicin aged fibroblasts without fixation. After cells were harvested, they were washed with PBS and PBS containing 0.1% Tween 20. Nuclei were released with TM2 (20 mM Tris–HCl, pH: 8.0; 2 mM MgCl2) with 0.5% Nonidet P 40 Substitute (Sigma Cat #74385). Afterward, nuclei were washed with TM2, and chromatin was either digested for 4 min for nChIP or 8 min for ChIP-seq with 1.0 U MNase (Sigma Cat #N3755-500UN) in nuclei solubilized with 1 mL of 0.1 M TE (10 mM Tris, 0.2 mM EDTA, 100 mM NaCl) and supplemented with 1.5 mM CaCl2. MNase reactions were quenched with 10 mM EGTA and centrifuged at 1000 rpm at 4 °C. Supernatant was removed, and nuclei extracted overnight at 4 °C in 0.35M/0.6M buffer supplemented with a protease inhibitor cocktail (Roche Cat #05056489001). ChIP was performed with anti-H3K4me2/ anti-H3K9me3 antibody. nChIP’ed chromatin bound to Protein G Sepharose beads (GE Healthcare Cat #17-0618-02) were washed 3X with cold 0.5X PBS. Following this, DNA was extracted from the chromatin bound Protein G-Sepharose beads followed by qRT PCR.

### Chromatin fractionation

Nuclei from both young and aged fibroblasts were isolated and the cell pellet was then treated with Micrococcal nuclease (MNase) (2 U/ml) for 3-4 min, and chromatin was extracted overnight from each nuclei using a low-salt buffer, and 0.35MNaCl containing buffer with 1× Complete Protease Inhibitor Cocktail (catalog no. 11697498001, Millipore Sigma, USA) at 4°C. The following day the chromatin fractions were collected after spinning down the nuclei pellet at 8000rpm for 10 mins. Pooled nuclei pellet from the above fractions was then further subjected to 0.6M NaCl containing buffer for 4 hours. All the chromatin fractions were collected in 2 biological replicates from young and aged cells and subjected to mass spectrometry.

### RNA extraction and quality control

RNAs were extracted by TRIzol reagent (catalog no. 15596026, Ambion, USA) according to the manufacturer’s protocol. One T175 flask of cells was resuspended in 1 ml of TRIzol and incubated at room temperature for 5 min. The cell debris was spun down at 12,000 rpm for 10 min at 4°C, and the supernatant was transferred to a fresh tube. For 1 ml of TRIzol, 200 μl of chloroform was added and incubated for 2 min at room temperature. After centrifugation at 12,000 rpm for 15 min at 4°C, the clear phase was transferred to a fresh tube and mixed with 500 μl of isopropanol (catalog no. 534021, Sigma-Aldrich). The samples were centrifuged again, and the RNA pellet was washed with 1 ml of cold 75% ethanol and resuspended in diethyl pyrocarbonate ultrapure water (KD Medical, USA). The RNA samples were subjected to DNase I treatment to avoid genomic DNA contamination using a DNase I kit [New England Biolabs (NEB), USA] following the manufacturer’s protocol. The integrity of RNA was verified by resolving in 1% agarose gel containing GelStar nucleic acid gel stain (catalog no. 50535, Lonza, USA) in a Mupid-One gel electrophoresis system (TaKaRa, Japan). RNA samples were purified for the second time by the TRIzol method as mentioned above and were stored at −80°C until further analysis.

### Complementary DNA synthesis

After quantifying the RNAs using a DeNovix DS-11 spectrophotometer (DeNovix Inc., USA), 1 μg of RNA was taken to synthesize complementary DNA (cDNA) conversion reactions were done using an iScript™ Advanced cDNA Synthesis Kit (catalog no. 18080085, Biorad, USA) following the manufacturer’s protocol. The cDNAs were diluted in nuclease-free water and stored at −20°C until use.

### Quantitative real-time polymerase chain reaction

cDNA samples were prepared using the GreenLink™ Blue qPCR Mastermix (catalog no. 16-2200, BioLink laboratories, USA) following the manufacturer’s protocol. The quantitative reverse transcription PCR (qRT-PCR) was run on the StepOnePlus Real-time PCR System (Applied Biosystems, USA), and relative quantification was performed using the 2^−ΔΔCT^ method. All the real-time PCRs were performed in 20-μl volume and triplicates. Control reactions without the template were performed to rule out nonspecific amplification (primer dimers). GAPDH and Actin served as internal controls respectively. Melting curve analysis was performed for all the primer sets to check the specificity of the primers. Primer sequences used in this study are listed in the table.

### Data base analysis of transcriptomic data

The GTEx version 8 RNA-seq data set was obtained from the GTEx Portal website (https://storage.googleapis.com/adult-gtex/bulk-gex/v8/rnaseq/GTEx_Analysis_2017-06-05_v8_RNASeQCv1.1.9_gene_tpm.gct.gz).

Heatmaps were downloaded using the R package TCGA biolinks. In total, we analyzed 3,061 RNA-seq data from six tissues (heart, lung, liver, skin, colon, and kidney) provided by GTEx, groups (young: 20-39, middle-aged: 40-59, and old: 60-79). Specifically, the GTEx dataset included 626 samples from the heart, 385 from the lung, 158 from the liver, 1258 from the skin, and 64 from the kidney. The GTEx samples were distributed as follows: 446 from the young group, 1,338 from the middle-aged group, and 1,277 from the old group.

Five groups of genes were selected for the heatmap: Inner Kinetochore (CENPN, CENPL, CENPO, CENPQ, CENPU, CENPH, CENPI, CENPK, CENPM, CENPS, CENPT, CENPW, CENPX), Centromere (CENPA, CENPC, CENPB), Outer Kinetochore (MIS12, DSN1, NSL1, PMF1, NDC80,NUF2, KNL1, SPC24, SPC25, ZWINT), Chaperons (DAXX, HIRA, HJURP), and Others (CENPE, CENPF, CENPJ, CENPV). For the heatmap, FPKM values were calculated from the count data and Actin gene expression was used to normalize the expression across the samples. Genes were then sorted within each group based their categories.

### RNA Sequencing

For RNA expression profiling, total RNA was isolated from ∼. Sequencing libraries were constructed from total RNA samples with the Illumina TruSeq Stranded Total RNA Library Prep Kit (RS-122-2201). Libraries were multiplexed and sequenced on the NextSeq instrument using NextSeq High 2.1 chemistry and sequenced for 151 cycles in paired-end mode. Reads of the samples were trimmed for adapters and low-quality bases using Cutadapt (v1.18) before alignment with the human reference genome (hg38) with STAR/RSEM (v 2.7.0f) tools. The following parameters were used: —alignIntronMin20 — sjdbScore 1 —peOverlapNbasesMin10. To calculate differential expression, read counts within genes were generated with Subreads v.2.01, against the Ensembl release 103 annotations. Differential expression was calculated with DESeq and is reported at a significance threshold of false discovery rate–adjusted P value of <0.01. Transcript abundance estimates were reported in units of RPKM (reads per kilobase of transcript per million reads mapped).

### Mass spectrometry

Experiment carried out independently twice for 2 biological replicates. Increasing amount of salt concentration was used to extract the chromatin fractions, 0.5X PBS represents Euchromatin fractions whereas 0.35M and 0.6M NaCl fractions are the heterochromatin fractions. Protein concentrations were measured using Bicinchoninic Acid protein Assay (BCA). From these concentrations, 25ug of each sample were taken for digestion. Proteins were denatured using Rapigest, reduced with DTT, and then Alkylated with IAA. An SP3 digestion was performed using a magnetic bead handling robot to wash beads prior to trypsin digestion. Samples were incubated overnight at 37 degree C in trypsin. Rapigest was hydrolyzed by quenching with acid.

Mass Spectrometry. Peptides were analyzed with an Evosep One coupled with a Thermo Exploris 480 tandem mass spectrometer. For each injection, we loaded approximately 1 μg peptides on Evosep tips and separated them using an 88-minute separating gradient. For data independent acquisition (DIA) analysis, we performed chromatogram library experiments as described in ^35,53^. Briefly, we acquired 6 chromatogram library acquisitions with 4 m/z DIA isolation windows using a staggered window pattern from narrow mass ranges. For all single-injection acquisitions, the Thermo Exploris 480 was configured to acquire 75 × 8 m/z (covering 400-1,000 m/z) precursor isolation windows using an optimized staggered window pattern^54^. MS1 full scans were interspersed every 75 MS/MS spectra. Isolation window schemes have been previously described ^53^.

### Preparation of metaphase chromosomes

Colcemid (catalog no. 10295892001, Millipore Sigma, USA) was added for 12-16 hours in young/aged fibroblasts, and the cells were harvested by trypsinization (catalog no. 25200056, Thermo Fisher Scientific, USA). The cells were washed with 10 ml of 1× PBS, suspended in 6 ml of hypotonic solution, and incubated in a water bath at 37°C for 20 min. They were fixed using freshly prepared cold fixative solution (methanol: glacial acetic acid in 3:1 proportion). They were suspended in 4 ml of fixative following centrifugation at 1500 rpm for 5 min at room temperature and incubated for 1 hour at room temperature. They were again centrifuged and resuspended in 400 μl of the fixative solution and stored at 4°C until slide preparation.

About 40 to 80 μl (two to three drops) of the fixed cells were dropped over the cold glass slide prewashed with fixative and were air-dried for 3 min. The slides were viewed under a light microscope to check the density of the metaphase spread, and the same protocol was repeated with cell dilutions using a freshly prepared fixative solution until optimal metaphase spread and density were attained. The slides were immediately processed for immunofluorescence.

### Cell cycle synchronization of BJ skin Fibroblasts

For cell-cycle-dependent analysis in mitosis, both young and aged cells were arrested by double-thymidine treatment. Cells were first seeded onto coverslips for immunofluorescence. Cells were blocked with a final concentration of 5 mM thymidine for 18 hr. Cells were released and grown in fresh media for 9 hr, followed by a second cell-cycle block with 5 mM thymidine for 12 hr. Synchronized cells were released into M phase by washing and cultivation in fresh media for another 10-11 hours. Mitotic phases were identified by DAPI staining of DNA, which was stimulated with 405 Diode laser at low intensity and detected via a 420 nm long path filter.

### Immunofluorescence

Cells were grown on poly-D-lysine coated coverslips. The media was removed, and the cells rinsed with cold 1X PBS. This was followed by prefixing the cells on the coverslip with 1% PFA in PEM (80 mM K-PIPES (pH: 6.8), 5 mM EGTA (pH: 7.0), 2 mM MgCl2) for 10 min. After washing the cells with cold PEM, it was permeabilized using 0.5% Triton-X in CSK (10 mM K-PIPES (pH: 6.8), 100 mM NaCl, 300 mM sucrose, 3 mM MgCl_2_, 1 mM EGTA) for 5 min at 4°C. 6. The coverslips were then rinsed with PEM and fixed with a few drops of 4% PFA in PEM at 4°C for 20 min. This was followed by a wash with PEM and then permeabilization with 0.5% Triton-X in PEM at room temperature for 5 minutes. The cells were then blocked using PBS + 3% BSA + 5% normal goat serum/10 uL/mL normal goat IgG for 1 hr at room temperature (or overnight at 4°C). This was followed by incubation with the primary antibody to desired dilution in hybridization solution for 2-3 hours. Following washes with 1× PBS supplemented with 0.05% Tween, slides were incubated for 1 hr at room temperature with Alexa Fluor 488/568/647 goat anti-IgG (1:1500, Invitrogen). DNA was counterstained with DAPI (50 ng/ml in 1× PBS). The coverslips were observed with a DeltaVision RT system (Applied Precision) controlling an interline charge-coupled device camera (Coolsnap; Roper) mounted on an inverted microscope (IX-70; Olympus). Images were captured by using a 100× objective at 0.1 μm z sections, deconvolved, and projected by using softWoRx. Three biological replicates were performed for each experiment.

For co-immunofluorescence both BJ Young and doxorubicin induced aged cells were transfected with 2ug of HA-LSD-1/Flag-CENP-B plasmid (Sinobiologicals). After 72 hours transfected cells grown on coverslips were fixed and co immunostained as above using anti-mouse -Flag antibody and anti-HA -Rabbit antibody. Colocalization of proteins on interphase nuclei was identified using the “Colocalization” plugin in ImageJ software. Colocalization readings from each young and aged^dox^ nuclei were calculated and the percentage was calculated for total number of CENP-B/ HA-LSD-1 foci. Intensity from the IF image was measured using ImageJ and normalized against the background signal.

### Immunoblotting

The cell was lysed in radioimmunoprecipitation assay (RIPA) buffer supplemented with 1× protease inhibitor cocktail and centrifuged for 10 min at 10,000 rpm at 4°C. The supernatant was collected and mixed with 4× Laemmli sample buffer (catalog no. 1610747, Bio-Rad, USA), then denatured for 7 min at 95°C and incubated on ice for 2 min. The samples were loaded into 4 to 20% Mini-PROTEAN TGX precast gels (catalog no. 4561093, Bio-Rad, USA), resolved in 1× tris-glycine SDS running buffer (#RGC-3390, KD Medical, USA), and transferred to nitrocellulose membrane using Trans-Blot Turbo Mini Transfer Packs (catalog no. 1704158, Bio-Rad, USA). The membrane was blocked in 1:1 Odyssey blocking buffer (catalog no. 92740000, LI-COR, USA) and 1× phosphate-buffered saline (PBS) at room temperature for 1 hour and incubated with primary antibody diluted in 1:1 blocking buffer and 1× PBS complemented with 0.1% Tween 20 on a rocker for overnight at 4°C. After three washes in 1× PBS, 0.1% Tween 20, the membrane was incubated with the Alexa Fluor conjugated to the secondary antibody diluted in blocking buffer complemented with 0.1% Tween 20% for 1 hour at room temperature. As mentioned previously, the membrane was washed and imaged in an Odyssey CLx scanner (LI-COR Biosciences, USA) and analyzed using Image Studio Lite (LI-COR Biosciences, USA).

All antibodies used for immunoblotting and immunofluorescence in this study are commercially available and are listed in the key resource table.

### Human Tissue harvesting and processing

Tissues of the lungs and kidneys of three different age groups were obtained from Amsbio. The details of the tissue are included in the table. Frozen tissue weighing 0.4-0.5g was cut into two equal fractions and processed for protein and RNA isolation. For nuclear preparation, around 0.2-0.3g of tissue was suspended in Buffer A (250mM sucrose, 15mM TrisHcl, pH 8.0, 15mM NaCl, 60 mM KCl, 1 mM EDTA, 0.5 mM EGTA, 1 mM Spermidine, Protease inhibitor) for 15 minutes on ice followed by a wash with buffer A. The tissues were dounced with a homogenizer on ice no more than 30 seconds. The dounced samples were then centrifuge for 10 min at 4°C at 1,500 rpm.

The pellet was resuspended again in 10 mL fresh Buffer A and centrifuged subsequently for 10 min at 4°C at 1,500 rpm. The buffer was removed and proceeded to Nuclear Prep/Extraction or RNA isolation through trizol method.

### Statistical analysis

All the numerical data are presented with SEM in graphs. The intensity of each centromeric foci was counted for each age/young group in biological triplicates. The intensity the IF signals was presented with p values. The differences between means from the signal intensities were analyzed by two-tailed Student’s t test (Mann-Whitney) with Welch’s correction. All statistical differences were calculated using GraphPad Prism software (v7.7e, GraphPad Software Inc., USA). A P value of <0.05 was considered as a statistically significant difference. Illustrative diagrams were created using BioRender (BioRender, USA) and Photoshop CC 2019 (Adobe Inc., USA).

## Acknowledgments

The authors acknowledge the important feedback received from A Ganesan, D Melters, M Bui, J Hsia, A Gilbert for critical feedback and advice on the work. We thank CCR Core Sequencing Facility for sequencing, total RNA. We thank K McKinon from NCI CCR Vaccine Branch Flow Cytometry Core for the flow cytometry analysis. We thank D. Melters and P. Sen (NIA) for assistance in chromatin extraction protocol. We acknowledge T. Misteli (CCR/NCI) for all the intellectual inputs and advice for this work. We also thank B. Earnshaw for sharing the CENP-C antibody which was used extensively in this study.

## Funding

This work was supported by the Intramural Research Program of the NCI, NIH to Y.D.

## Author contributions

S.S. and Y.D. conceptualized the research, analyzed the data, and wrote the manuscript. S.S. designed and performed the experiments and acquired and analyzed the data. S.B. analyzed the total RNA sequencing data and developed karyoplots. T.M performed cell synchronization and mitotic assays and analyzed independently the mitotic defects.

## Competing interests

The authors declare that they have no competing interests.

## Limitations of the study

In this manuscript, we demonstrate for the first time that the essential centromeric proteins CENP-A and CENP-C decline in both senescent and aged human fibroblasts. We also report downregulation of centromeric transcription and centromeric proteins in aged tissues from various origins (kidney and lung). However, human aging is a complex biological process governed by multiple genetic and epigenetic factors. Consequently, one limitation to our work is that the degree of downregulation observed in centromeric proteins during older age may differ depending on the specific tissue or cellular origin. Another caveat is there is an inherent challenge in tracking the same individual across multiple timepoints throughout the lifespan in a longitudinal fashion. Our approach to rejuvenate mitosis in aged cells using a lysine demethylase inhibitor requires validation in an organismal model. Chromatin-regulatory complex inhibitors likely have additional targets outside the centromeres, which will be the focus of follow up work. Lastly, our current data indicates that CENP-B colocalizes with LSD-1 in aged cells, potentially leading to the suppression of H3K4me2 at the centromeric DNA. However, due to the lack of a suitable LSD-1 antibody, we are unable to unequivocally demonstrate a physical interaction of LSD-1 with the centromeric protein CENP-B, which will require development of further tools.

## Notes

### Competing Interest Statement

The authors have declared no competing interest.

### Summary of Updates

This manuscript has been revised for updated figures

