## Supplemental Figures for "Centromere inactivation during aging can be rescued in human cells"

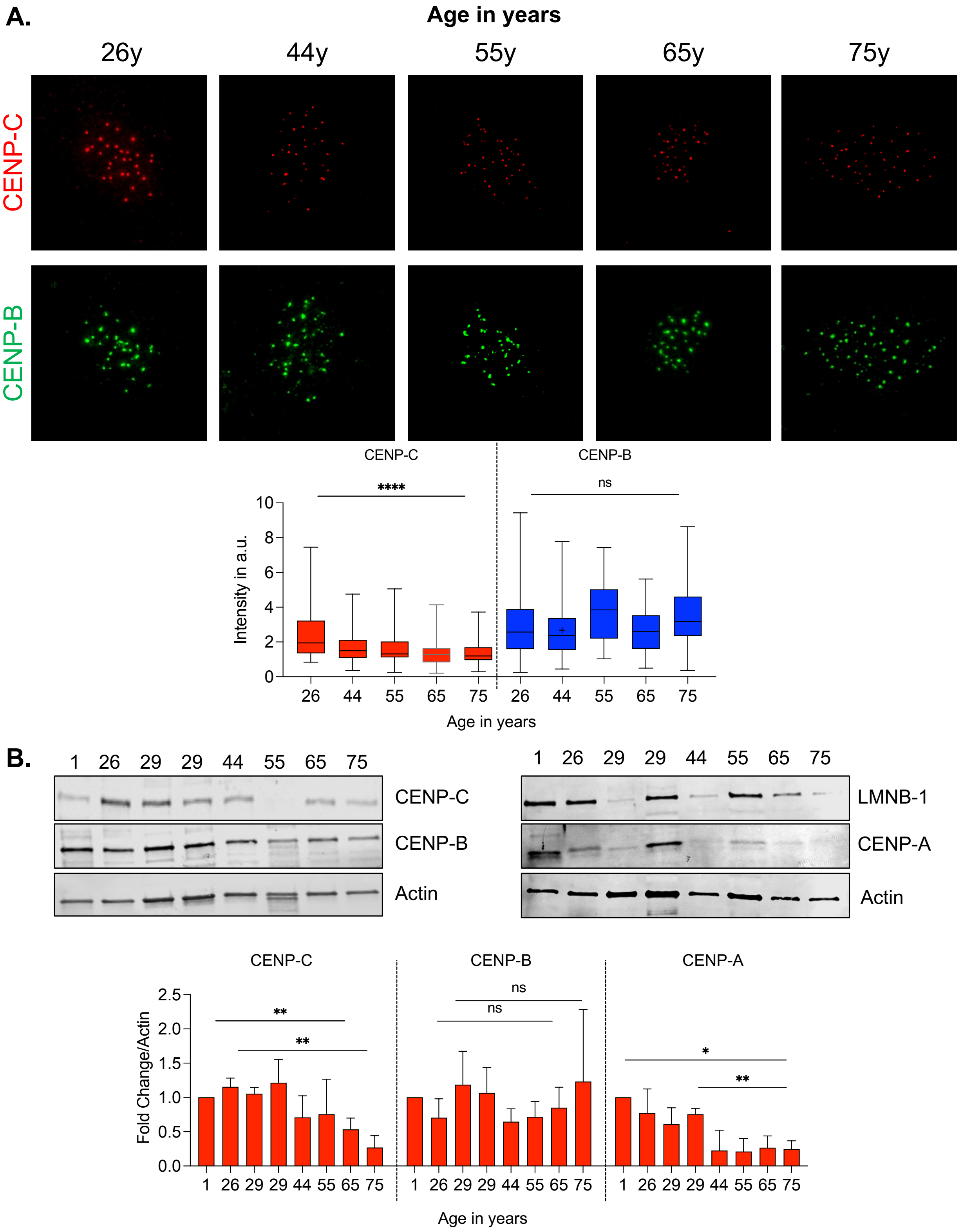


**Figure S1:** (A) Representative IF image of CENP-C (in red) and CENP-B (in green) levels in different aged skin fibroblasts. No.of nuclei used for quantification denoted by n for each age is mentioned as follows: 26y (n=52), 44y (n=37), 55y (n=38), 65y (n=38), 75y (n=57). Scale bars as denoted. (B) Quantitation of fluorescence intensity of CENP-C (red) and CENP-B (green) across different aged skin fibroblasts. CENP-C intensity diminishes in age 75y as compared to the young fibroblasts of age 26 y. CENP-B levels however remain unaltered. Representative immunoblot images of proteins isolated from different aged fibroblasts. CENP-A/C levels are reduced in aged cells, but no significant difference is observed for CENP-B. Quantifications of immunoblots from 3 different biological replicates are represented in bar graphs. All data are shown as means ± SEM. **P < 0.001 and ****P < 0.0001.

A.

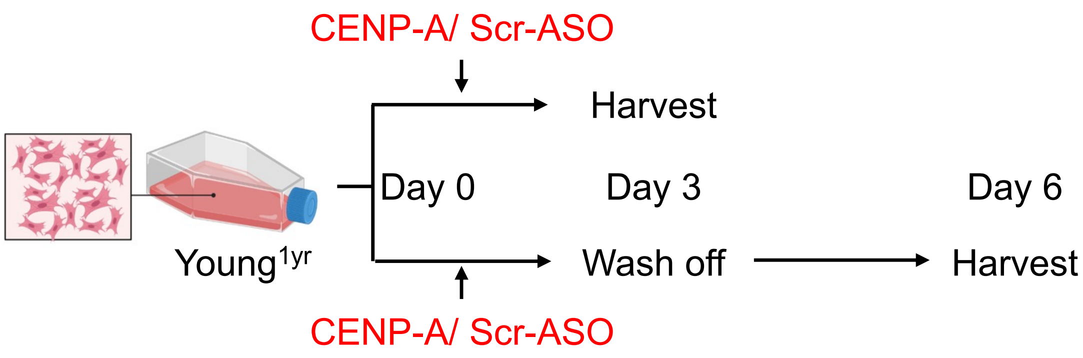


B.

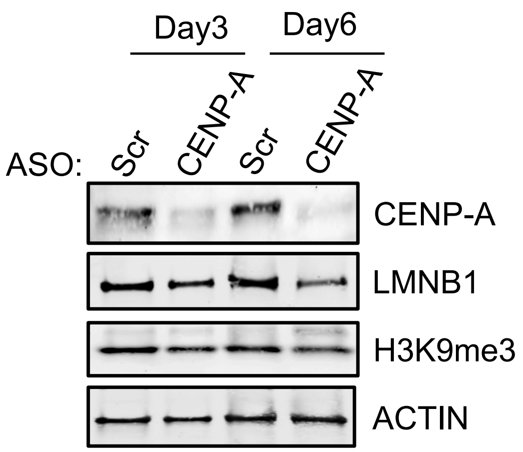


**Figure S2:** (A) Scheme depicting the transfection of antisense oligonucleotides specific to CENP-A in young 1 year old skin fibroblasts. (B) Decrease in CENP-A level lead to the reduction of Lamin B1 levels. Significant reduction in levels of lamin B1 observed post 6 days of transfection.

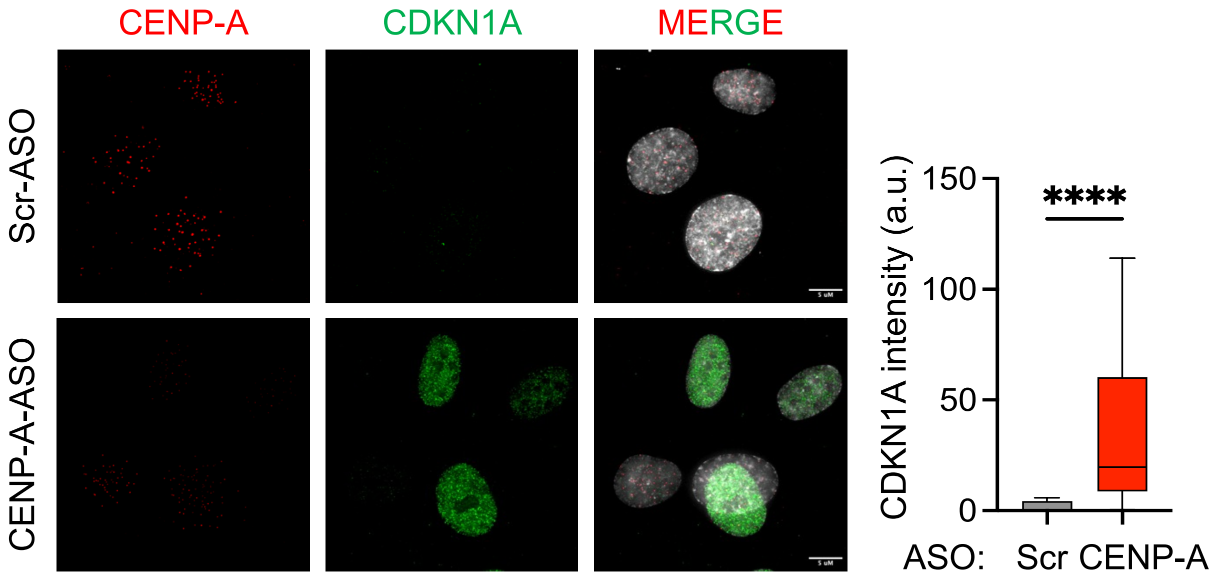


**Figure S3:** Immunofluorescence depicting CDKN1A expression upon CENP-A knockdown post 72 hours in 1 year old fibroblasts. Knockdown of CENP-A induces CDKN1A mediated cellular senescence. Data represented from n>40

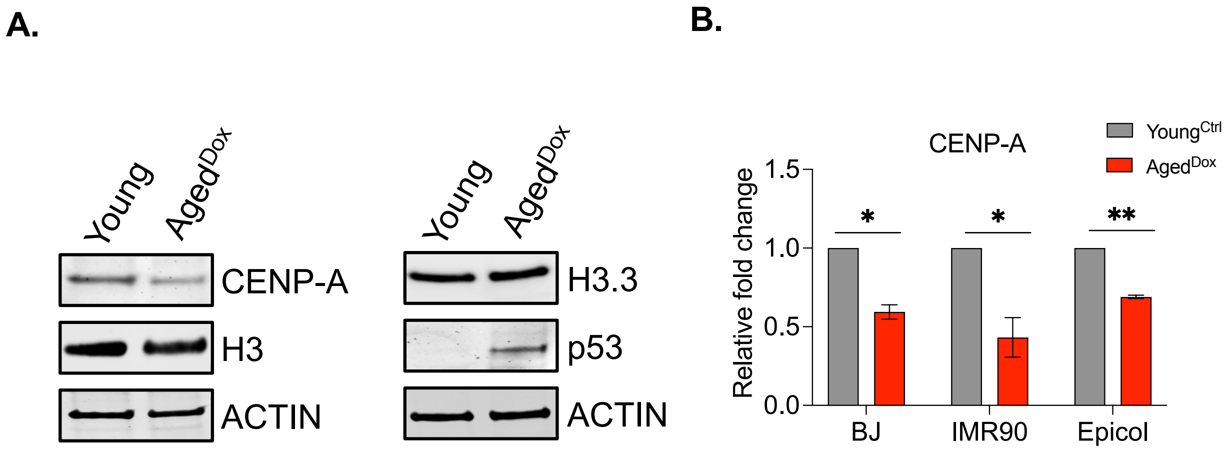


**Figure S4:** (A) Representatives immunoblot images of proteins isolated from BJ DMSO-treated young control cells and doxorubicin-induced senescent cells. CENP-A levels are reduced in senescent cells, but no significant reduction is observed for histone H3.3. p53 protein was found to be upregulated specifically in the senescent cells. (B) Quantitation of immunoblotting of CENP-A from young and doxorubicin induced senescent cells of various origin BJ (skin fibroblast), IMR90 (lung fibroblast), Epicol (Colon epithelial cell line). Data from 3 biological replicates. All data are shown as means ± SEM. **p < 0.001 and ****p < 0.0001.

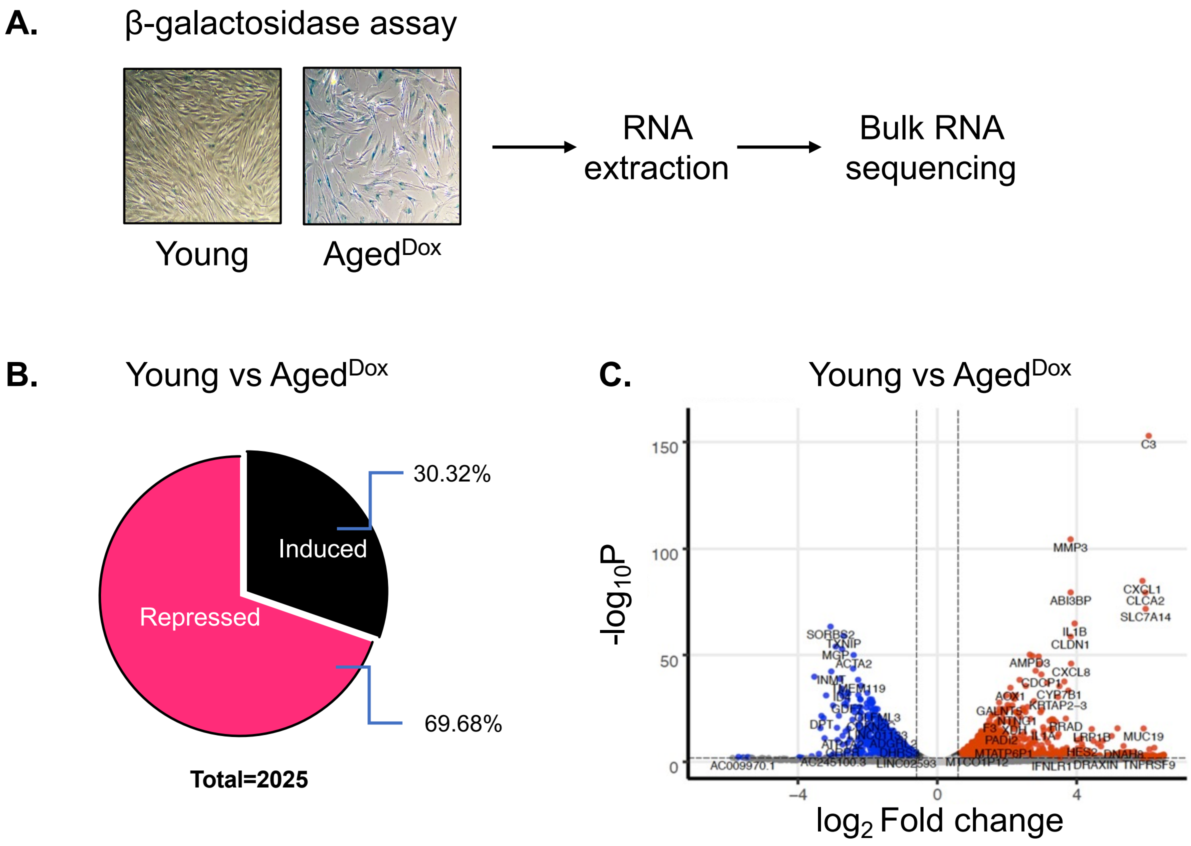


**Figure S5:** (A) Beta galactosidase assay of BJ skin fibroblasts Young (DMSO control) and Doxorubicin induced senescent cells. Both cells subjected to bulk RNA sequencing. (B) Vein diagram signifying the total number of genes repressed and induced in doxorubicin senescent cells vs young cells. (C) Volcano plot depicting differentially expressed genes. Red signifies significantly upregulated genes while blue denotes downregulated genes. Data represented from 2 biological replicates.

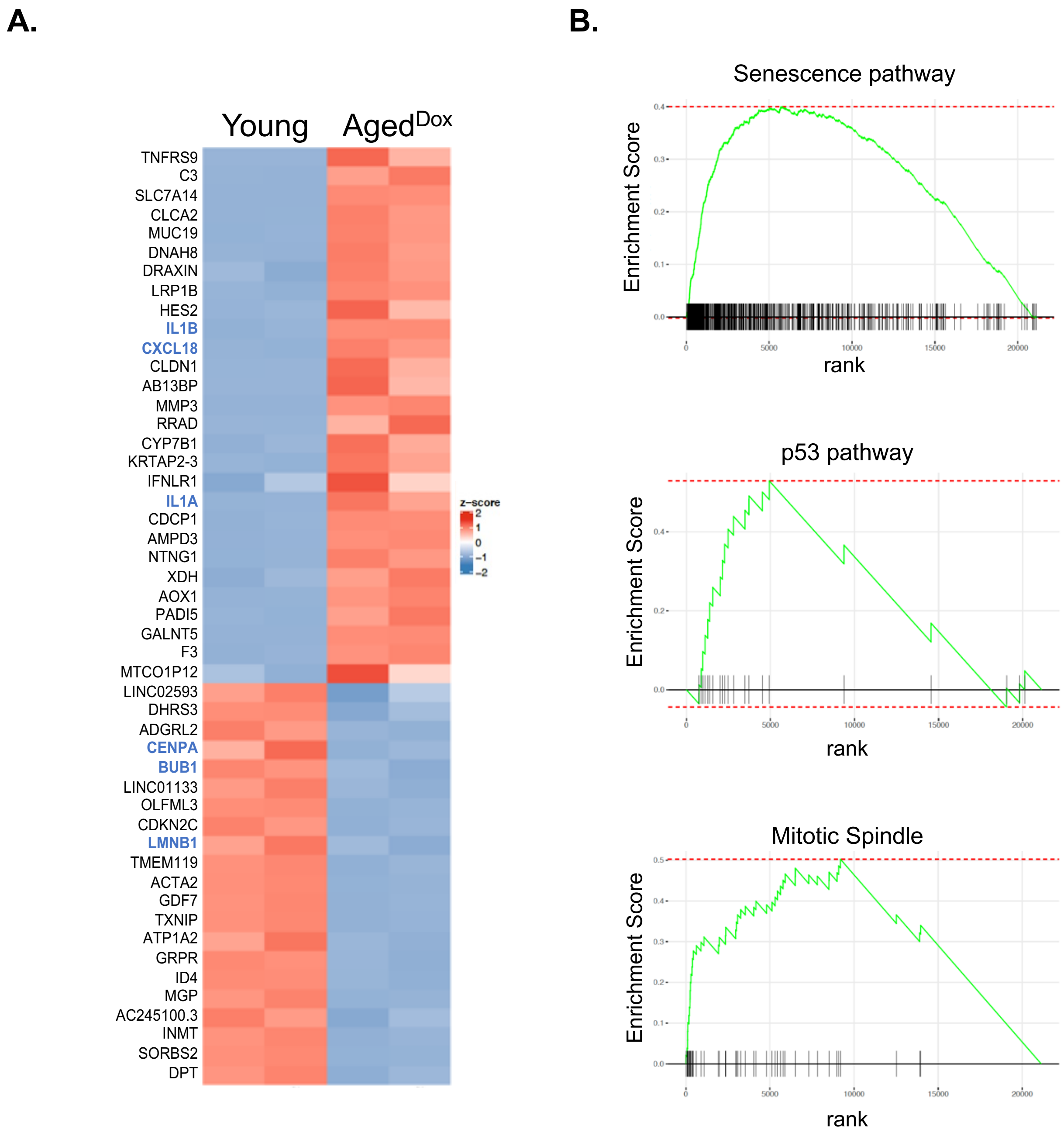


**Figure S6:** A. Heat map depicting top 50 differentially expressed genes in young versus Aged$^{Dox}$ cells. Red signifies significantly upregulated genes while blue denotes downregulated genes. Genes of interest denoted in blue. B. Enrichment plots of pathways perturbed upon aging.

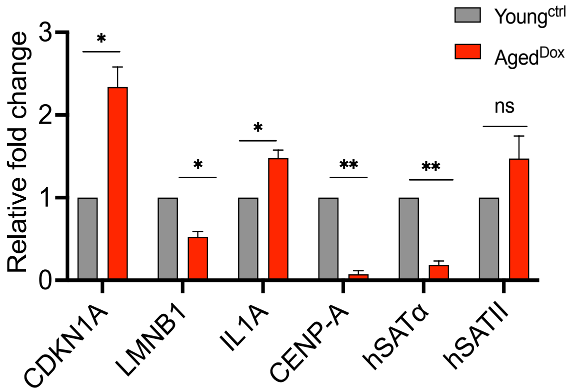


**Figure S7:** qRT PCR results showing RNA expression of different genes in young vs doxorubicin aged cells. Both CENP-A and centromeric transcript hSATα RNA was reduced in BJ skin doxorubicin aged fibroblasts. Data represented from 3 biological replicates.

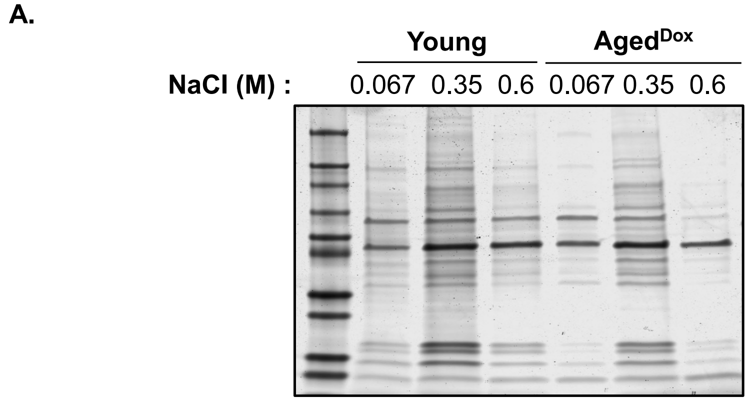


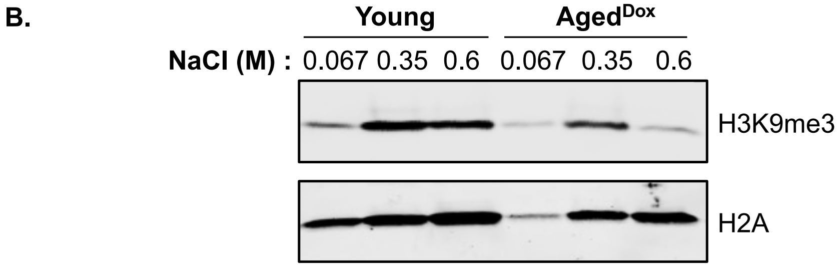


**Figure S8:** (A) Gel profile of chromatin fractions extracted with increasing salt concentration from young and doxorubicin aged BJ skin fibroblasts. (B) Immunoblot analysis of heterochromatin mark H3K9me3 reveal differential expression pattern across different salt fractions in young vs aged BJ skin fibroblasts.

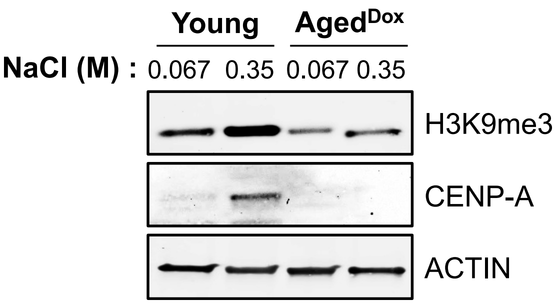


**Figure S9:** Immunoblot analysis of heterochromatin mark H3K9me3 reveal differential expression pattern across different salt fractions in young vs aged BJ skin fibroblasts. CENP-A expression decreases in aged cells. In young cells CENP-A is present majorly in the heterochromatin fraction.

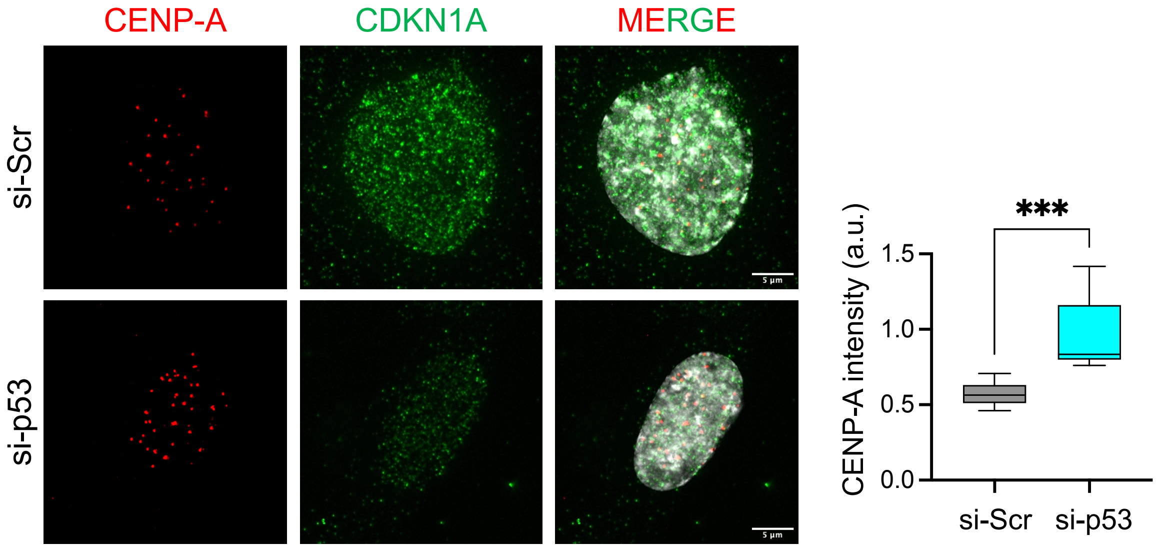


**Figure S10:** Immunofluorescence depicting CDKN1A expression upon p53 knockdown in 47year old skin fibroblasts. Knockdown of p53 induces expression of CENP-A in 47-year-old skin fibroblasts.

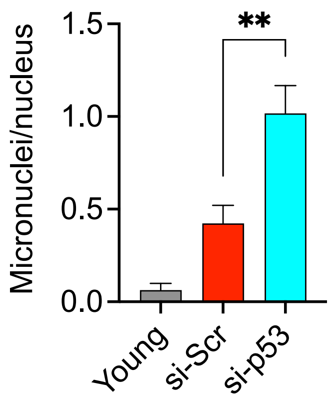


**Figure S11:** Quantitation of micronuclei in Young versus aged and in aged p53 knockdown cells. Significant accumulation of micronuclei was observed in both aged and p53 knockdown aged cells. Data represented n>50.

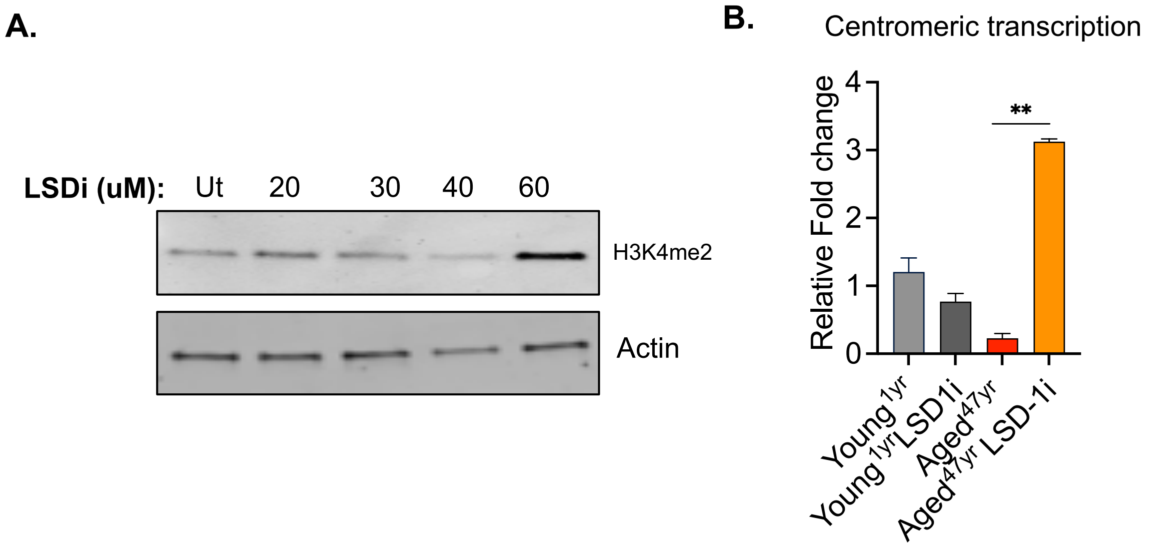


**Figure S12:** (A) Representative immunoblot image to show H3K4me2 levels upon LSD-1 inhibition at different concentrations in BJ aged$^{dox}$ cells. (B) qRT PCR for centromeric transcription upon 2-PCPA LSD-1 inhibition in 1year vs 47-year-old skin fibroblasts. Data represented from 3 biological replicates.

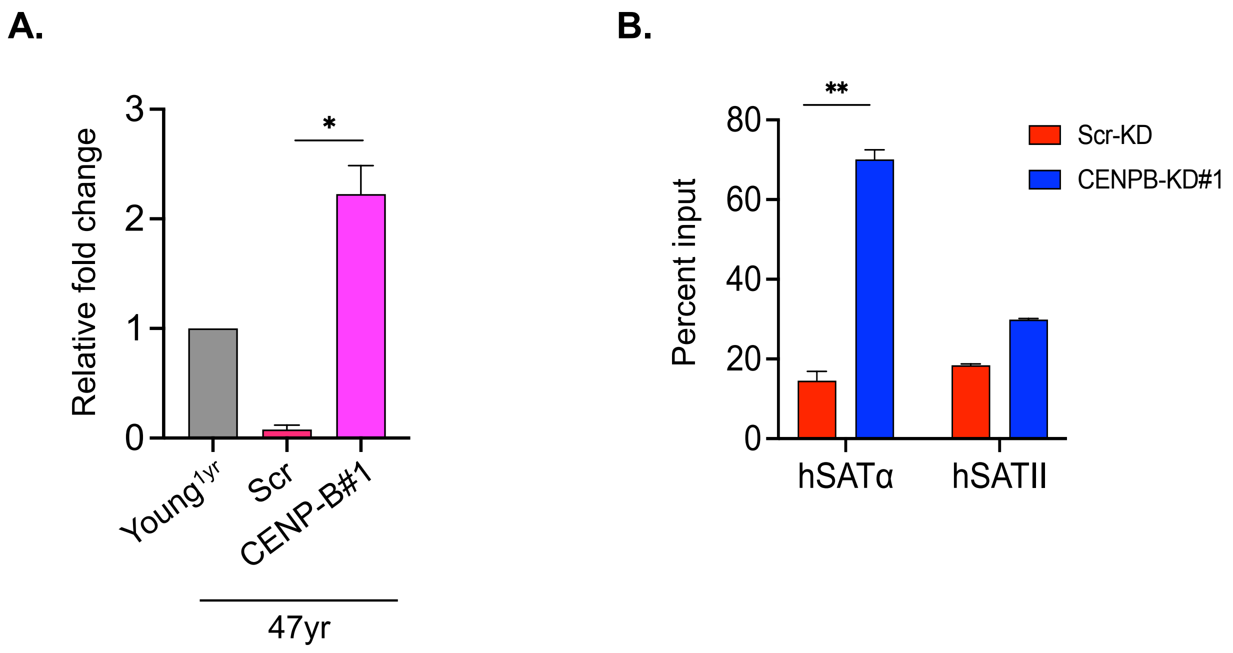


**Figure S13:** (A) qRT PCR for centromeric transcription upon CENP-B knockdown in 47-year-old skin fibroblast post 72 hours of transfection. Data represented from 3 biological replicates. (B) Relative enrichment of H3K4me2 at the centromeric as well as pericentromeric DNA upon CENP-B knockdown in doxorubicin induced aged cells. Data represented from 2 biological replicates

Quantitation tables given below:

Figure 1B

| Fibroblast | N (number of Nuclei) | Mean CENP-C Intensity | *p*-value |
|---|---|---|---|
| AG08498 1y | 49 | 1.984 | |
| AG09860 26y | 92 | 2.353 | |
| AG04441 29y | 53 | 3.226 | |
| AG13156 44y | 82 | 1.323 | <0.0001 (29 vs 44) |
| AG06858 47y | 53 | 0.7986 | |
| AG06283 55y | 56 | 1.698 | |
| AG04659 65y | 78 | 1.167 | <0.0001(26 vs 65) |
| AG06883 75y | 58 | 1.027 | <0.0001(26 vs 75) |
| GM00731 96y | 33 | 0.7442 | |

Figure1C

| Fibroblast | N (number of foci) | Mean CENP-C Intensity | *p*-value | Mean CENP-B Intensity | *p*-value |
|---|---|---|---|---|---|
| AG06103A 29y | 122 | 2.063 | | 1.861 | |
| AG13156 44y | 150 | 0.9076 | <0.0001 | 1.771 | 0.4865 |
| AG04659 65y | 105 | 1.136 | <0.0001 | 2.281 | 0.0191 |

Figure 2B:

| BJ Fibroblast | Mean CENP-C intensity | N ((number of Nuclei) | *p*- value |
|---|---|---|---|
| Young | 1.823 | 81 | |
| Aged$^{>PD}$ | 1.274 | 50 | 0.0012 |
| Aged$^{Dox}$ | 0.5262 | 79 | <0.0001 |

Figure 2C:

| BJ Fibroblast | Mean $\gamma$-H2AX intensity | N ((number of Nuclei) | *p*- value |
|---|---|---|---|
| Young | 2.441 | 141 | |
| Aged$^{>PD}$ | 5.438 | 67 | <0.0001 (Aged$^{>PD}$ vs Young) |
| Aged$^{Dox}$ | 4.265 | 77 | 0.5066 (Aged$^{>PD}$ vs Aged$^{Dox}$) |

Figure 4C:

| BJ Fibroblast (Aged$^{Dox}$) | Mean CENP-C intensity | N (number of Nuclei) | *p*- value |
|---|---|---|---|
| si-Scr | 1.138 | 76 | |
| si-p53 | 2.523 | 79 | <0.0001 |

Figure 4D:

| BJ Fibroblast | Mitotic defect (fraction) | N (number of Nuclei) | *p*- value |
|---|---|---|---|
| Young | 0.04545 | 61 | |
| Aged$^{Dox}$ si-Scr | 0.2689 | 93 | 0.0103 |
| Aged$^{Dox}$ si-p53 | 0.2320 | 142 | 0.0040 |

Figure 6B:

| BJ Fibroblast (Aged$^{Dox}$ si-p53) | % Bridge | % Lagging | N (number of nuclei) | No. of mitotic cells |
|---|---|---|---|---|
| -LSD-1i | 50.00 | 12.5 | 145 | 44 |
| +LSD-1i | 5.26 | 7.89 | 87 | 37 |

Figure 6C:

| BJ Fibroblast (Aged$^{Dox}$ si-p53) | Mean Fraction of Micronuclei/Nuclei | N (number of nuclei) | *p-value* |
|---|---|---|---|
| -LSD-1i | 1.088 | 225 | |
| +LSD-1i | 0.4209 | 167 | <0.0001 |

Figure 7B:

| BJ Fibroblast | N (number of foci) | Mean CENP-C Intensity | *p*-value | Mean CENP-B Intensity | *p*-value |
|---|---|---|---|---|---|
| Young | 51 | 5.429 | | 5.145 | |
| Aged$^{Dox}$ | 39 | 2.296 | <0.0001 | 4.686 | 0.3231 |

Figure S1A:

| Fibroblast | N (number of foci) | Mean CENP-C Intensity | *p*-value | Mean CENP-B Intensity | *p*-value |
|---|---|---|---|---|---|
| AG09860 26y | 52 | 2.504 | | 2.942 | |
| AG13156 44y | 34 | 1.734 | 0.0101 | 2.677 | 0.4881 |
| AG06283 55y | 38 | 1.595 | 0.0023 | 3.778 | 0.0453 |
| AG04659 65y | 38 | 1.479 | 0.0003 | 2.718 | 0.5237 |
| AG06883 75y | 57 | 1.463 | <0.0001 | 3.574 | 0.0792 |

Figure S3:

| AG08498 1y Fiborblast | Mean CDKN1A intensity | N (number of Nuclei) | *p*- value |
|---|---|---|---|
| Scr-ASO | 4.348 | 29 | |
| CENP-A-ASO | 36.44 | 40 | <0.0001 |

Figure S10:

| AG06858 47y Fiborblast | Mean CENP-A intensity | N (number of Nuclei) | *p*- value |
|---|---|---|---|
| Scr-ASO | 0.5372 | 19 | |
| CENP-A-ASO | 2.130 | 30 | 0.0003 |

Figure S11:

| BJ Fibroblast | Mean Fraction of Micronuclei/Nuclei | N (number of Nuclei) | *p*- value |
|---|---|---|---|
| Young | 0.03509 | 139 | |
| Aged$^{Dox}$ si-Scr | 0.9186 | 132 | <0.0001 |
| Aged$^{Dox}$ si-p53 | 1.088 | 225 | <0.0001 |
