## Supplemental Table for "Centromere inactivation during aging can be rescued in human cells"

| **Primer Name** | **Primer sequence (5’—3’)** |
| --- | --- |
| hSATII Fp | AATCATCGAATGGTCTCGAT |
| hSATII Rp | ATAATTCCATTCGATTCCAC |
| IL1A Fp | TGTATGTGACTGCCCAAGATGAAG |
| IL1A Rp | AGAGGAGGTTGGTCTCACTACC |
| LMNB1 Fp | GAAAAAGACAACTCTCGTCGCA |
| LMNB1 Rp | GTAAGCACTGATTTCCATGTCCA |
| b-ACTIN Fp | AGATGTGGATCAGCAAGCAG |
| b-ACTIN Rp | TCCTCGGCCACATTGTGAACTTTG |
| hSATα Fp | CAAGAAGGCTTCAAAGCACC |
| hSATα Rp | TTCATTGGAATCGCGAATAC |
| GAPDH Fp | TCGACAGTCAGCCGCATCT |
| GAPDH Rp | CTAGCCTCCCGGGTTTCTCT |
| hSATII ChIP Fp | CATCGAATGGAAATGAAAGGAGTC |
| hSATII ChIP Rp | ACCATTGGATGATTGCAGTCAA |

Table S1: List of Primers used in the study.

| **Sample ID** | **Gender** | **Age** | **Ethnicity** | **Sample Type** | **Tissue Type** | **Diagnosis** | **PMI, hours** | **BMI** | **Cause of death/Reason of Surgery** |
| --- | --- | --- | --- | --- | --- | --- | --- | --- | --- |
| 041449B1(1) | Male | 1 | Caucasian | Formalin Fixed | lung | normal tissue | N/A | N/A | Cystic malformation |
| 046015A (4) | Male | 60 | Caucasian | Formalin Fixed | lung | normal tissue | 4.5 | 26.0 | open craniocerebral injury, multiple trauma of bones and internal organs after train accident |
| 044487A (5) | Male | 88 | Caucasian | FF | lung | normal tissue | 2.5 | 27.7 | post MI |
| 051829A (2) | Male | 23 | Caucasian | FF | kidney | normal tissue | 3 | 37.9 | neck phlegmon, PE |
| 053041A(2) | M | 60 | Caucasian | FF | kidney | normal tissue | 4.5 | 26.0 | open craniocerebral injury, multiple trauma of bones and internal organs after train accident |
| 05735A(2) | M | 80 | Caucasian | FF | kidney | normal tissue | 2.5 | N/A | ischemic stroke, cerebral edema, brain stem dislocation |

Table S2: Tissue details used in the study.
